# Rhesus Cytomegalovirus-encoded Fcγ-binding glycoproteins facilitate viral evasion from IgG-mediated humoral immunity

**DOI:** 10.1101/2024.02.27.582371

**Authors:** Claire E. Otero, Sophia Petkova, Martin Ebermann, Husam Taher, Nessy John, Katja Hoffmann, Angel Davalos, Matilda J. Moström, Roxanne M Gilbride, Courtney R. Papen, Aaron Barber-Axthelm, Elizabeth A. Scheef, Richard Barfield, Lesli M. Sprehe, Savannah Kendall, Tabitha D. Manuel, Nathan H. Vande Burgt, Cliburn Chan, Michael Denton, Zachary J. Streblow, Daniel N. Streblow, Scott G Hansen, Amitinder Kaur, Sallie Permar, Klaus Früh, Hartmut Hengel, Daniel Malouli, Philipp Kolb

**Author notes:** Co-corresponding authors: Daniel Malouli, Vaccine and Gene Therapy Institute, Oregon Health and Science University, 505 NW 185th Ave., Beaverton, OR 97006., Philipp Kolb, Institute of Virology, Medical Center, Faculty of Medicine, University of Freiburg, Hermann-Herder-Str. 11, 79104 Freiburg, Germany. These authors contributed equally to this work (Claire E. Otero, Sophia Petkova, Martin Ebermann).

## Abstract

Human cytomegalovirus (HCMV) encodes four viral Fc-gamma receptors (vFcγRs) that counteract antibody-mediated activation *in vitro*, but their role in infection and pathogenesis is unknown. To examine the *in vivo* function of vFcγRs in animal hosts closely related to humans, we identified and characterized vFcγRs encoded by rhesus CMV (RhCMV). We demonstrate that Rh05, Rh152/151 and Rh173 represent the complete set of RhCMV vFcγRs, each displaying functional similarities to their respective HCMV orthologs with respect to antagonizing host FcγR activation *in vitro*. When RhCMV-naïve rhesus macaques were infected with vFcγR-deleted RhCMV, peak plasma viremia levels and anti-RhCMV antibody responses were comparable to wildtype infections. However, the duration of plasma viremia was significantly shortened in immunocompetent, but not in CD4+ T cell-depleted animals. Since vFcγRs were not required for superinfection, we conclude that vFcγRs delay control by virus-specific adaptive immune responses, particularly antibodies, during primary infection.

## Introduction

Cytomegaloviruses (CMVs) are a group of widespread, strictly species-specific β- herpesviruses found in various mammals including primates (*1, 2*). Primary infection of CMV- naïve, immunocompetent hosts generally resolves asymptomatically followed by lifelong persistent infection with periodical reactivation events (*3*). Repeated exposure to viral antigens results in extraordinarily strong CMV-specific humoral and cellular immune responses that control, but do not clear viral infection (*3, 4*). In immunocompromised individuals such as transplant recipients, uncontrolled human CMV (HCMV) infections can lead to severe morbidity and mortality (*5*). Furthermore, in congenitally infected neonates HCMV is the leading cause of non-genetic birth defects including sensorineural hearing loss (*6*) and neurological deficits (*7*). Thus, the development of a prophylactic HCMV vaccine for vulnerable populations has been designated a “Tier I priority” by the Institute of Medicine since 2000 (*8*). To date however, all HCMV vaccine approaches have failed in phase II clinical trials (*9–11*).

As productive HCMV infections are restricted to humans, we previously established a congenital CMV (cCMV) infection model in rhesus macaques (RMs) which allows for the examination of exploratory treatment and vaccine approaches in an evolutionarily closely related host (*12*). In this model, we demonstrated that pre-exposure administration of highly concentrated IgG purified from plasma of RMs with potent anti-CMV IgG responses (HIG) can limit the replication and congenital transmission of rhesus CMV (RhCMV) (*13*). Although no consistent benefit of post-exposure HIG therapy has been demonstrated for cCMV infections across various human clinical trials, it appears that treatment dosage and pharmacokinetics are important parameters of efficacy and protection (*14–18*). In line with these observations, we recently demonstrated that not neutralization, but IgG-Fc-gamma (Fcγ)-mediated anti-viral mechanisms by non-neutralizing IgG, specifically maternal plasma levels of antibody-dependent cellular phagocytosis (ADCP) and antibody-dependent cellular cytotoxicity (ADCC), correlate with a reduced risk of congenital infection (*19, 20*). Intriguingly, both ADCP and ADCC are mediated by Fcγ receptors (FcγRs) expressed by nearly all immune cells including neutrophils, macrophages and NK cells.

Several structurally unrelated viral IgG Fc-binding glycoproteins (vFcγRs) have been identified for different herpesviruses: the gE/gI glycoprotein complex encoded by herpes simplex virus (HSV)-1 genes *US8* and *US7* respectively (*21–23*), gE encoded by Varicella zoster virus (VZV) (*24–26*) as well as various glycoproteins examined across different CMV species (*27–32*). The HCMV genome encodes four vFcγRs, *RL11* (RL11 or gp34), *UL119/118* (UL119/118 or gp68), *RL12* (RL12 or gp95) and *RL13* (RL13 or gpRL13) (*29, 33*). While these proteins can independently interfere with host FcγR activation *in vitro*, UL119/118 and RL11 can also act cooperatively (*34, 35*).

We recently identified Rh05 (*RL11A*) as the first vFcγR encoded by RhCMV and demonstrated that Rh05 interferes with host FcγR activation *in vitro.* However, Rh05-deleted RhCMV did not display any defect *in vivo*, possibly due to the expression of additional vFcγRs (*28*). Here, we identify Rh152/151 *(Rh152/151)* and Rh173 (*Rh173*) as additional vFcγRs encoded by RhCMV. Deletion of all three vFcγRs from a previously described full-length (FL) RhCMV clone representative of low passage primary isolates (*2*), resulted in restoration of antibody-mediated host FcγR activation *in vitro* and significantly shortened plasma viremia in immunologically intact, but not CD4+ T cell-depleted, CMV-naïve RM. These results establish CMV vFcγRs as *bona fide* immune evasion proteins and suggest that targeting vFcγRs might improve the effectiveness of anti-CMV vaccines or immunotherapies.

## Results

### *Rh152/151* encodes a UL119/118 vFcγR ortholog that can block host FcγR function

Since HCMV encodes multiple vFcγRs, we hypothesized that RhCMV should similarly encode vFcγRs in addition to Rh05 (*28*). As the RhCMV open reading frame (ORF) *Rh152/151* represents a direct sequence ortholog of the known HCMV vFcγR UL119/118 (gp68), we examined the corresponding protein for its IgG binding activity (*36*). The *Rh152/151* ORF in the fibroblast-adapted laboratory strain 68-1 contains a premature termination codon removing two YXXΦ sorting motifs (*2, 36*) (Fig 1A). Since C-terminal modifications of UL119/118 increased surface expression by decreasing internalization (*34*), we compared IgG-binding of intact Rh152/151, obtained from FL-RhCMV (*2*), with truncated, strain 68-1-derived Rh152/151 and Rh152/151 in which the predicted transmembrane and cytoplasmic domains were replaced with that of human CD4 (Fig 1A). All three Rh152/151 constructs were able to bind IgG, with both the truncated and CD4-tail version showing elevated IgG binding similar to the previously described CD4-tail version of HCMV UL119/118 (*34*). Similarly, when co-expressed with rhesus CD4 using a T2A linker sequence (*37*) to test their effect on FcγR activation by a rhesus CD4-specific rhesusized IgG, the truncated or CD4-tailed versions of Rh152/151 showed stronger antagonization of human CD16 activation compared to Rh152/151 from FL-RhCMV (Fig. 1B). Furthermore, similar to the effect of UL119/118 on human CD16 (*34*), Rh152/151 was able to directly interfere with the binding of rhesus CD16 to immune complexes formed by anti-CD20 IgG (rituximab) on cells expressing both CD20 and Rh152/151 (Fig. 1C). In contrast, Rh05 was unable to inhibit CD16 binding (Fig. 1C), consistent with previous results obtained for HCMV RL11 (gp34) (*34*). This suggests that RhCMV Rh152/151 represents a sequence and functional ortholog of HCMV UL119/118 (gp68).

**Fig 1.**
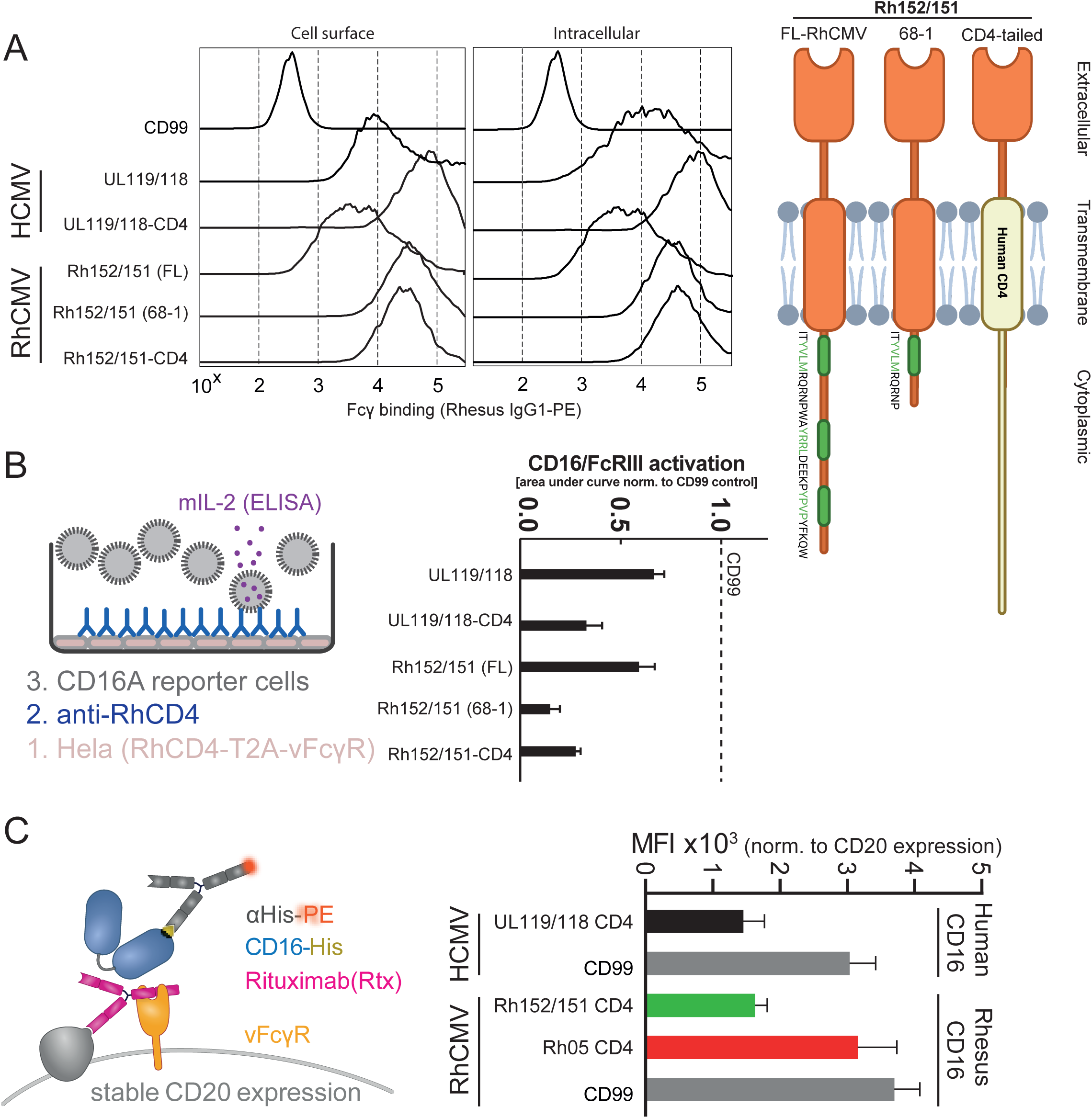
Rh152/Rh151 binds IgG and antagonizes host FcγR activation by blocking receptor engagement. A) 293T cells were transfected with pIRES-eGFP plasmid expressing the indicated UL119/118 or Rh152/Rh151 constructs or human CD99 as a negative control. Polycistronic GFP expression was used to gate on transfected cells. The cells were probed for binding of PE-conjugated rhesusized IgG1 by flow cytometry with (intracellular) or without permeabilization (cell surface). The cartoon illustrates the orientation and configuration of the different Rh152/Rh151 constructs in the cellular membrane. Extracellular and cytoplasmic termini of the proteins are indicated and the modifications to the YXXΦ sorting motifs are highlighted in green. B) Transfected Hela cells expressing the RhCD4 target antigen and the indicated vFcγRs in equimolar amounts from a T2A-linked fusion protein were incubated with graded amounts of RhCD4-specific rhesusized IgG1 and tested for human CD16 activation using a cell-based reporter assay as described before (*34*). The bar graph shows area under curve (AUC) values of the response curves of three independent experiments performed in technical replicates and normalized to activation in the presence of a non-Fcγ binding glycoprotein control (human CD99). Error bars = +SD. C) 293T cells stably expressing human CD20 were transfected with the indicated vFcγRs or human CD99, incubated with the anti-CD20 antibody Rituximab and probed for rhesus or human CD16 binding via flow cytometry. The experimental setup is shown in detail on the left. The bar graphs on the right show the MFI normalized to CD20 expression following transfection across three independent experiments. Error bars = +SD.

### The highly polymorphic *RL11* family member Rh173 (*RL11T*) encodes for an IgG binding protein that shows conserved antagonization of host Fc**γ**R activation

In HCMV, three out of four identified vFcγRs are members of the *RL11* gene family, a class of highly diverse glycoproteins clustered near the 5’ terminus of the genome. In order to determine whether the *RL11* family members encoded by RhCMV contain vFcγRs in addition to Rh05, we inserted every *RL11* family member annotated in the RhCMV genome into a p-IRES-eGFP vector. To monitor surface expression, we replaced the predicted N-terminal signal peptides with the signal peptide of human tapasin followed by a hemagglutinin (HA) tag. Upon transfection of 293T cells, we measured rhesus IgG1 binding via flow cytometry (Fig 2A). HA-staining demonstrated that transient transfection of plasmids containing ORFs Rh06 (*RL11B*), Rh08 (*RL11D*), Rh08.1 (*RL11E*) and Rh22 (*RL11L*) showed very low expression levels (Fig. 2A). However, in their original sequence these genes lack either a signal peptide or a transmembrane domain making them unlikely to encode for vFcγR candidates. Of the *RL11* family members that do encode for both of these features only Rh19 and Rh23 demonstrated below average expression levels, yet expression was likely sufficient to detect Fcγ binding. As expected, most of the *RL11* family members did not display IgG binding either at the cell surface or intracellularly upon permeabilization of the transfected cells. The notable exceptions were Rh05 which we described previously and Rh173, but not Rh13.1, the orthologue of HCMV RL13 which was previously reported to bind human IgG (*33*) (Fig 2A). The *Rh173* ORF is the only *RL11* family member not located near the 5’ end of the genome but in a central region homologous to the *ULb’* region of HCMV. Rh173 shows substantial amino acid sequence diversity across RhCMV isolates, a feature it shares with the HCMV vFcγR RL12 (gp95) (*38*) (Fig. S1). To determine whether these sequence variants can antagonize host FcγR activation equally well, we tested several Rh173 genotypes from different RhCMV isolates in a cell-based FcγR activation reporter assay (*28, 39*) (Fig. 2B). Plasmids containing the *Rh173* ORFs were co-expressed with rhesus CD4 via a T2A linker sequence and transfected HeLa cells were incubated with a rhesus CD4-specific rhesusized mAb and human CD16 reporter cells. All examined Rh173 sequence variants showed a similar reduction of CD16 activation, suggesting a conserved function not affected by the polymorphisms in the Rh173 ectodomains. In conclusion, we have identified orthologs for all four known HCMV vFcγRs, but, since Rh13.1, the ortholog of HCMV RL13, did not show any IgG binding, we conclude that RhCMV encodes three functional vFcγRs.

**Fig 2.**
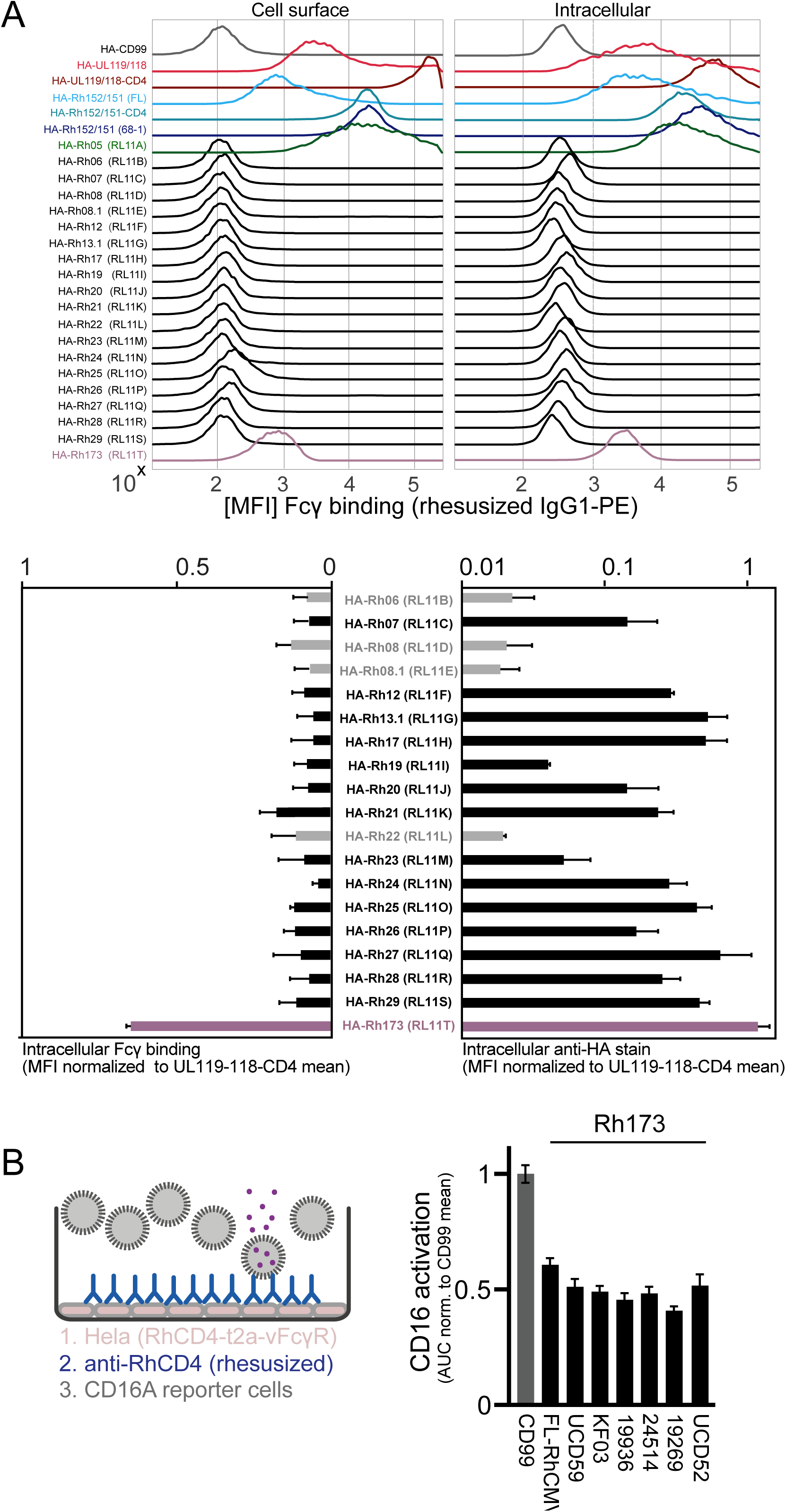
Rh173 binds IgG and antagonizes FcγR activation. A) 293T cells transfected with a pIRES-eGFP plasmids providing a Tapasin signal peptide and an N-terminal HA-tag encoding either a known vFcγR, a RhCMV *RL11* family member or the human CD99 protein as a negative control were probed for binding of PE-conjugated rhesusized IgG1 with (cell surface) or without permeabilization (intracellular) via flow cytometry. The presented histograms show a representative experiment. The bar graphs depict the MFI (from two independent experiments) for intracellular IgG-binding (left panels) or intracellular protein expression using the N-terminal HA-tag (right panels) and normalized to UL119/118-CD4 results. Error bars = range. The samples presented with grey bars show RL11 family members that lack a predicted signal peptide and/or transmembrane domain in their original sequence. B) Hela cells were transfected with plasmid pIRES-eGFP expressing a T2A-linked fusion protein of the RhCD4 target antigen and the indicated Rh173 sequence variants from published RhCMV sequences. The transfectants were incubated with graded amounts of RhCD4-specific rhesusized IgG1 and tested for human CD16 activation using a cell-based FcγR activation reporter assay. Equal transfection efficiency was monitored via polycistronic GFP expression. The presented bar graph shows the area under the curve (AUC) values of the response curves of three independent experiments and normalized to activation in the presence of a non-Fcγ binding glycoprotein control (CD99). Error bars = +SD.

### RhCMV vFc**γ**Rs are expressed with similar kinetics, localize to subcellular vesicular structures and are integrated into newly formed virions

To characterize the RhCMV vFcγRs in the context of infection, we constructed a FL-RhCMV recombinant devoid of all three vFcγR. We additionally replaced the *Rh13.1* ORF with an SIVgag antigen which has the dual effect of stabilizing the virus genome during in vitro propagation and allowing us to specifically monitor immune responses to heterologous antigens as described before (*2, 40*) (Fig. 3A). Using FL-RhCMVΔRh13.1/SIVgag as parental recombinant we generated the triple-deletion mutant by sequentially deleting *Rh05, Rh152/151* and *Rh173*. As expected, the corresponding mRNAs were no longer detectable in infected fibroblasts whereas expression of ORFs flanking the deletions was unaffected (Fig. 3B). The deletion of all three vFcγRs also did not affect virus replication in rhesus fibroblasts (RFs) when compared to FL-RhCMV (Fig. 3C). Similarly, vFcγR deletion did not alter the ability of FL-RhCMV to infect epithelial cells *in vitro*, which is in contrast to a FL-RhCMV-derived recombinant that lacks two subunits of a pentameric complex (PC) required for infection of most cell types other than fibroblasts (Fig. 3D).

**Fig. 3.**
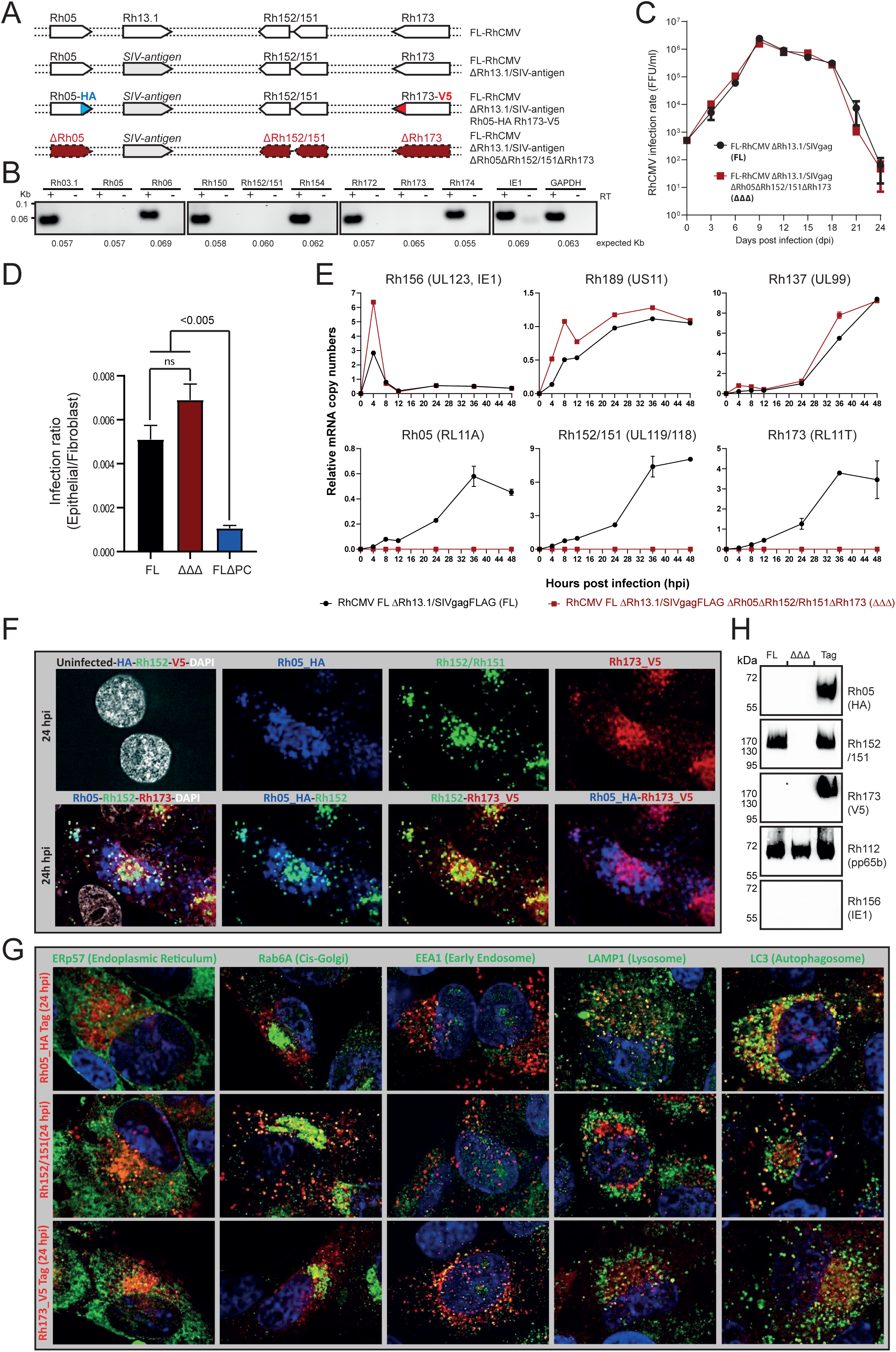
RhCMV vFcγRs are early late gene products that are non-essential for growth *in vitro* and locate to sub-cellular vesicular structures in infected cells. A) Graphical overview of FL-RhCMV-derived recombinants in which vFcγRs were either deleted or fused to HA or V5 epitope tags. HA and V5 epitope tags are highlighted in blue and red respectively. Deleted vFcγRs are indicated by highlighting the annotated ORFs in red. B) mRNA expression of vFcγR-encoding or neighboring genes was determined by RT PCR. Total mRNAs were harvested at 36h after infection of rhesus fibroblasts (RFs) with the triple vFcγRs deletion mutant (ΔΔΔ) (MOI=5). To determine mRNA expression cDNA was produced by reverse transcriptase (RT) and PCR was performed using primers targeting either the vFcγR-encoding or the neighboring ORFs. Control samples with no RT treatment were included. RhCMV IE1 and GAPDH were included as viral and cellular mRNA controls. C) A multistep growth-curve on RFs was performed to compare the replication kinetics of vFcγR-deleted recombinant (ΔΔΔ) compared to the vFcγR-intact parental FL-RhCMV-derived recombinant. RFs were infected at an MOI of 0.01, supernatants were collected at the indicated time points and the viral titers were determined as focus forming units (FFU) per ml of supernatant. The arithmetic mean of three biological repeat experiments are shown (± SEM). D) RFs and rhesus retinal pigment epithelial cells (RPEs) were infected with serial dilutions of FL-RhCMV, the triple vFcγR-deletion recombinant (ΔΔΔ), or a FL-RhCMV-derived recombinant deleted for the pentameric complex (PC) components Rh157.5 (UL128 homolog) and Rh157.4 (UL130 homolog). The infection rates in each cell type were determined by IFA for pp65b and are presented as the relative ratio of infected epithelial cells versus fibroblasts. E) The expression kinetics of the vFcγRs was monitored by infecting RFs with either FL-RhCMV or the triple vFcγR-deletion mutant (ΔΔΔ) at an MOI of 5. Total mRNA was harvested at 0, 4, 8, 12, 24, 36, and 48 hpi and cDNA was produced. The copy numbers of the viral transcripts for each gene determined by real-time quantitative reverse transcription PCR (qRT-PCR) are graphed in relation to copy numbers of the cellular GAPDH transcript. *Rh156* (UL123, IE1), *Rh189* (US11), and *Rh137* (UL99) (upper panels) were used as controls representative of IE, E and L kinetic classes during the viral replication cycle. F) To determine whether vFcγRs co-localize with each other we infected tRFs with FL-RhCMV/Rh05-HA/Rh173-V5 at a MOI of 1 for 24 h. Cells were fixed, permeabilized and stained with anti-HA, anti-Rh152/151 and anti-V5 antibodies for detecting vFcRs alone (top row) or in combination with each other (bottom row). Uninfected tRF were used as controls. Shown are representative single cell views. Uncropped images are provided in Fig. S4A. A parallel analysis at 48 hpi is shown in Fig. S4B and Fig. S4C. Similar results were obtained in four independent experiments. Fig. S5 shows secondary antibody control staining. G) To determine the subcellular localization of vFcγRs we infected RFs with FL-RhCMV/Rh05-HA/Rh173-V5 as in F) and co-stained with antibodies recognizing the cellular markers ERp57 (endoplasmic reticulum), Rab6A (Golgi), EEA1 (early endosome), LAMP1 (lysosome) and LC3 (autophagosome) for detecting specific organelles. Top row: co-staining with Rh05-HA, Middle Row: co-staining with Rh152/151. Bottom row: co-staining with Rh173-V5. The data presented are representative single cell views. Uncropped images are provided in Fig. S6 (bottom row). Uninfected cells are shown in Fig. S6 (top row). A parallel analysis performed at 48 hpi is shown in Fig. S7. Similar results were obtained in four independent experiments. H) Virions of FL-RhCMV, the triple vFcγR-deletion mutant (ΔΔΔ) or FL-RhCMV/Rh05-HA/Rh173-V5 were purified over a discontinuous Nycodenz gradient. Lysates were generated and immunoblots were performed for Rh05 (HA), Rh152/151 and Rh173 (V5) as well as pp65 and IE as positive and negative controls respectively.

To determine vFcγR expression kinetics, we examined vFcγR mRNA expression levels over 48 h upon infection of primary rhesus fibroblasts with FL-RhCMV at an MOI of 3. As representative immediate early (IE), early (E) and late (L) genes we also monitored the expression of the viral mRNAs *Rh156* (UL123, IE1), *Rh189* (US11) and *Rh137* (UL99) (*41*). Our results indicate that the three RhCMV encoded vFcγRs are expressed with early-late (E-L) kinetics (Fig.3E), matching expression kinetics of HCMV encoded vFcγRs (*29*).

To be able to determine the subcellular localization of the vFcγRs in the context of viral infection we generated a recombinant FL-RhCMV carrying an SIVgag transgene in the Rh13.1 locus in which the C-terminus of Rh05 was fused to the influenza HA epitope whereas the C-terminus of Rh173 was fused to the parainfluenza V5 epitope (Fig. 3A). Rh152/151 was not epitope tagged because we discovered that a mAb (*42*) originally identified in a screen of RhCMV-specific mouse hybridoma cell lines recognized Rh152/151 (Fig S2) as well as Cynomolgus CMV Cy152/151 (Fig. S3A) although it is unable to neutralize Rh152/151-mediated antagonization of host CD16 activation (Fig. S3B). We infected tRFs with the recombinant FL-RhCMVΔRh13.1/SIVgag/Rh05-HA/Rh173-V5 at an MOI of 1 and monitored the subcellular localization of each vFcγR by immunofluorescence analysis (IFA). At 24 hpi, all three vFcγR displayed a punctate vesicular staining pattern consistent with type I transmembrane glycoproteins locating to subcellular vesicular structures (Fig. 3F). However, co-staining of the vFcγRs indicated distinct localization of Rh05 whereas the localization of Rh152/151 and Rh173 seemed to overlap at least partially. Co-staining with markers for specific cellular compartments indicated that, in addition to early endosomes, Rh152/151 and Rh173 showed some co-localization with the Golgi-marker Rab6, whereas Rh05 partially co-localized with lysosomal and autophagosomal markers LAMP-1 and LC-3 (Fig. 3G, Fig. S6). These staining patterns are consistent with vFcγRs being endocytosed and ultimately degraded in lysosomes at early times of infection. However, at later stages of infection at 48 hpi, both Rh152/151 and Rh173 localized to perinuclear structures in the cell that co-stained with Rab6 and likely represent the viral assembly compartment (VAC) (Fig. S4, Fig S7).

The presence of vFcγRs in the VAC suggested the possibility that some of them might be incorporated into virions during viral assembly or envelopment which is consistent with mass spectrometry of strain 68-1 virions (*36*). To determine whether vFcγRs are indeed present in virions we gradient-purified virions from the supernatant of fibroblasts infected with FL-RhCMVΔRh13.1/SIVgag/Rh05-HA/Rh173-V5, FL-RhCMVΔRh13.1/SIVgag or triple deleted FL-RhCMVΔRh13.1/SIVgag/ΔΔΔ. Interestingly, each vFcγR was detected in immunoblots of purified virions of FL-RhCMVΔRh13.1/SIVgag/Rh05-HA/Rh173-V5 whereas they were all absent from FL-RhCMVΔRh13.1/SIVgag/ΔΔΔ (Fig. 3H). As Rh152-151 was detected with a mAb directly as described above, Rh152/151 was also detected in FL-RhCMVΔRh13.1/SIVgag. As they encode a transmembrane domain, vFcγRs of RhCMV are likely to be inserted into the virion envelope during assembly matching published results for HCMV (*43*).

### RhCMV encoded vFc**γ**Rs antagonize host Fc**γ**R activation *in vitro*

Our data suggest that RhCMV Rh152/151, Rh05 and Rh173 are orthologs of HCMV UL119/118 (gp68), RL11 (gp34) and RL12 (gp95) respectively (Fig. 4A). To directly test this relationship regarding their function, we performed a side by side comparison of these proteins with respect to their ability to inhibit human CD16 activation using an FcγR activation assay (Fig. 4B). The levels of inhibition confirmed the suggested orthologs and indicate functional conservation between the RhCMV and HCMV vFcγRs.

**Fig 4.**
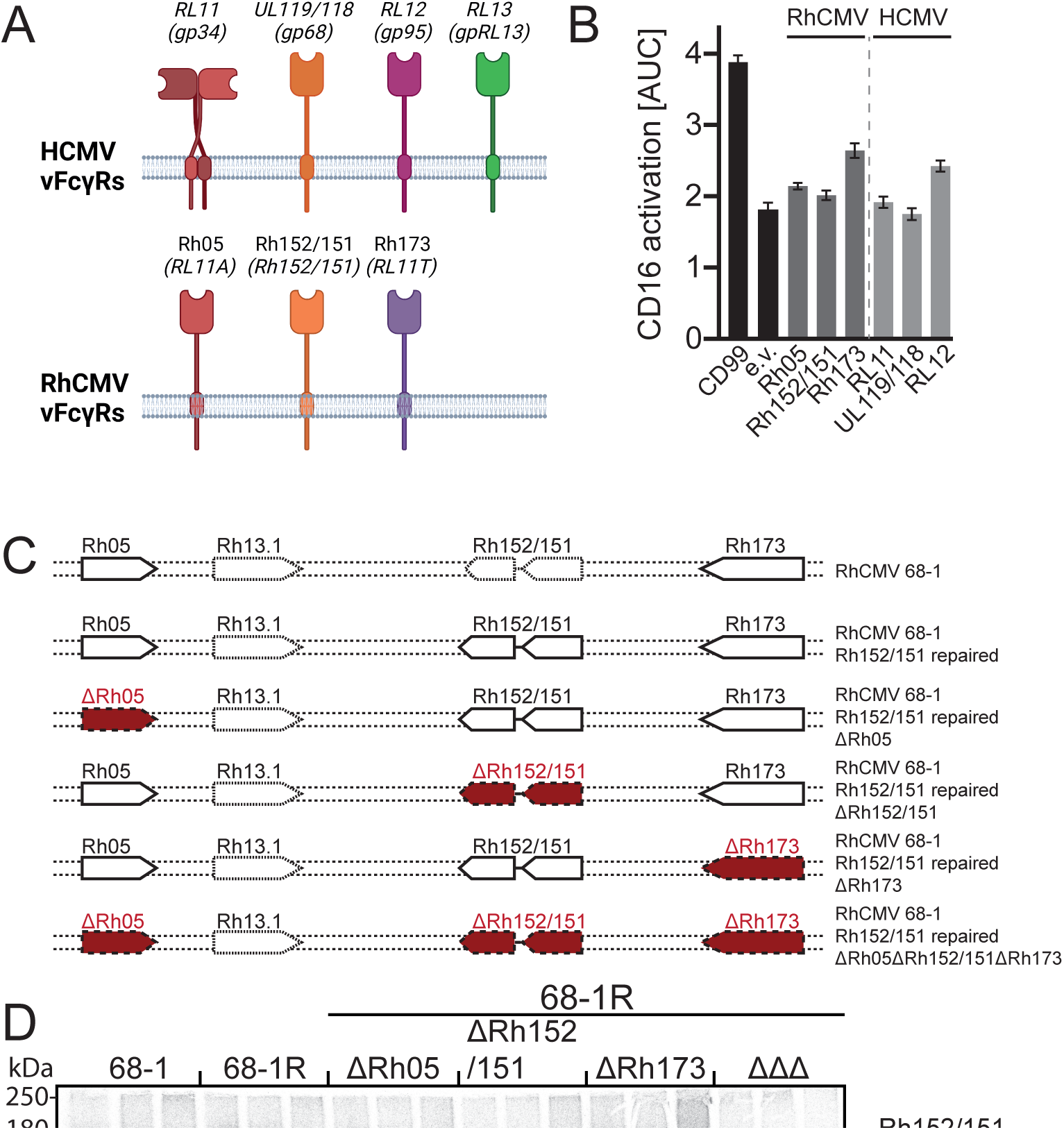
RhCMV vFcγRs reflect HCMV vFcγR function and efficiently antagonize host FcγR activation *in vitro*. A) Graphical overview of HCMV vFcγRs and their RhCMV orthologs. B) Transfected Hela cells expressing both the rhesus CD4 target antigen and the indicated vFcγRs as T2A-linked fusion protein from a pIRES-e-GFP plasmid were incubated with graded amounts of a rhesus CD4-specific rhesusized IgG1 antibody and tested for human CD16 activation using a cell-based reporter assay. Equal transfection was monitored via polycistronic GFP expression. The presented bar graph shows area under curve (AUC) values of the response curves of three independent experiments (Error bars = +SD). Empty vector (e.v.) transfection and expression of a non-Fcγ binding glycoprotein (CD99) served as controls. C) Overview over the RhCMV 68-1 based recombinants. To create RhCMV 68-1R, we repaired a premature termination codon in *Rh152/151*. Single and triple deletion mutants of the vFcγRs were based in the repaired 68-1 strain. All deletions are indicated by highlighting the corresponding ORFs in red. D) Telomerized rhesus fibroblasts (tRF) were infected with RhCMV 68-1 or 68-1R virus or 68-1R –derived single vFcγR deletion mutants at an MOI of 5 for 72 h and then labeled with [^35^S]-Met/Cys for 2 h. Whole cell lysates were prepared and precipitations of IgG binding molecules were performed using ProteinG bound rhesus-CD4 specific rhesusized IgG1. Lysates from each condition were split and either left untreated, or digested with either EndoH or PNGaseF. All samples were analyzed via gradient SDS-PAGE and subsequent autoradiography. One of two independent experiments is presented here. E) RFs were infected with RhCMV 68-1, 68-1R virus or single vFcγR deletion mutants at an MOI of 2 for 48h and then incubated with a 1:10 dilution of a pooled sera from eight CMV seropositive RM. The samples were assessed for human or rhesus FcγR activation using a cell-based reporter assay. Resulting mIL-2 ELISA OD values were normalized to the mean activation on vFcγR deleted virus for each subgroup. vFcγRs expressed by viruses used for infection are indicated below. *mock infected cells. Error bars = +SD. Three independent experiments performed in technical triplicates are presented here.

To determine the contribution of individual RhCMV vFcγRs to counteracting host FcγR activation by immune complexes we deleted each individual vFcγR or all three of them from a recombinant RhCMV strain 68-1 (68-1R) in which ORF *Rh152/151* was repaired by reversing the single nucleotide change resulting in a premature termination codon (Fig. 4C). Immunoprecipitations of IgG binding proteins using fully rhesusized IgG (anti-RhCD4, as above) from metabolically labeled tRFs infected with each of these recombinants revealed in each case that the expected number of vFcγRs were expressed by the respective recombinants whereas IgG binding proteins were not present upon deletion of all three vFcγRs (Fig. 4D). This result further supports the conclusion that we identified the full set of vFcγRs encoded by RhCMV. Furthermore, de-glycosylation using Endo H (cleavage of pre-Golgi N-glycans) or PNGase F (de-glycosylation of all N-glycans) confirmed RhCMV vFcγRs to be transmembrane glycoproteins that are processed via the ER-Golgi pathway similar to HCMV vFcγRs (*29*). Using these recombinant viruses, we further examined the ability of RhCMV vFcγRs to inhibit the activation of host FcγR in the context of viral infection. Infected RFs were incubated with plasma pooled from eight different RhCMV-seropositive RM followed by incubation with FcγR-activation reporter cells expressing single human or rhesus FcγRs associated with ADCP or ADCC. (Fig. 4E). Compared to RhCMV containing all three vFcγRs, an increased stimulation of all human and rhesus FcγRs was observed upon infection with RhCMV lacking all three vFcγRs consistent with them being able to antagonize all host FcγRs mediating ADCP or ADCC. This result also indicates that the FcγR activity is conserved between humans and RM consistent with previous IgG binding studies (*44*). The absence of either Rh05 or Rh152/151 alone resulted in a complete loss of antagonization whereas Rh173 deletion alone only marginally increased stimulation, which implies cooperation between Rh05 or Rh152/151 similar to HCMV UL119/118 and RL11 (*34*). Of note, the inhibitory effect on human and rhesus CD64, the only high affinity FcγR and main mediator of ADCP, was especially dependent on the presence of all three vFcγRs. These observations are consistent with previous observations regarding HCMV (*35*) and is in line with the synergistic mechanisms postulated between individual HCMV vFcγRs (*34*).

### A RhCMV mutant deleted for all three vFc**γ**Rs is cleared prematurely from circulation during primary infection in CMV naïve RM

To determine the role of vFcγRs on viral replication *in vivo*, we infected four RhCMV-naïve, male RM intravenously with 1×10^6^ PFU of the FL-RhCMVΔRh13.1/SIVgag recombinant deleted of all three vFcγRs (ΔΔΔ) and monitored the development of plasma viremia over time by qPCR. As controls, we infected three RhCMV-seronegative male RM with the parental FL-RhCMVΔRh13.1/SIVgag. We furthermore included historical results of 12 RhCMV-seronegative, female RM infected with RhCMVΔRh13.1/SIVgag into our control groups. Viral genome copy numbers became detectable in all animals shortly after inoculation, reaching peak levels around 7 dpi (Fig. 5A). However, the duration of the plasma viremia in animals infected with the vFcγR deletion mutant was significantly shortened compared to FL-RhCMV (median time to control 59.5 days versus 31.5 days post infection) (Fig.5B). The time point of viral clearance from plasma coincided with peak humoral responses of antibodies binding to whole virions and specific viral glycoproteins (gB, PC) (Fig. 5C), as well as antibodies mediating viral neutralization, ADCP or ADCC (Fig. 5D). Interestingly, the kinetics of the RhCMV-specific antibody responses were indistinguishable from RM inoculated with the parental recombinant. Together these results suggest that vFcγR-deletion does not affect the induction and specificity of the humoral immune responses, yet renders the virus more susceptible to antibody-mediated clearance. Intriguingly, when we depleted CD4 T-cells in RhCMV-naïve RM prior to inoculation we no longer observed any premature clearance of the (ΔΔΔ) deletion virus from the plasma in infected animals (Fig. 5E). Since CD4 T cell-depletion results in delayed RhCMV-specific antibody responses following inoculation (*12*), this result further supports a likely role of vFcγRs in counteracting the antibody response. Lastly, we investigated whether deletion of the vFcγRs would preclude superinfection of RhCMV seropositive RM that already have preexisting CMV specific adaptive immune responses. When inoculated two immunocompetent, RhCMV seropositive animals with FL-RhCMVΔΔΔ carrying a SIV-5’Pol antigen in the *Rh13.1* locus we observed strong and lasting SIV-5’Pol-specific CD4 and CD8 T-cell responses that developed shortly after inoculation, indicating that superinfection in the absence of humoral immune evasion genes is possible (Fig. 5F). In contrast, we previously reported that RhCMV recombinants devoid of viral gene products that inhibit MHC class-I mediated antigen presentation to CD8+ T cells lose their ability to superinfect RhCMV seropositive RM (*40*). Taken together, these results show that vFcγRs support primary infection of CMV-naïve hosts by counteracting antibody-mediated clearance from circulation, but they are not required for superinfection.

**Fig 5.**
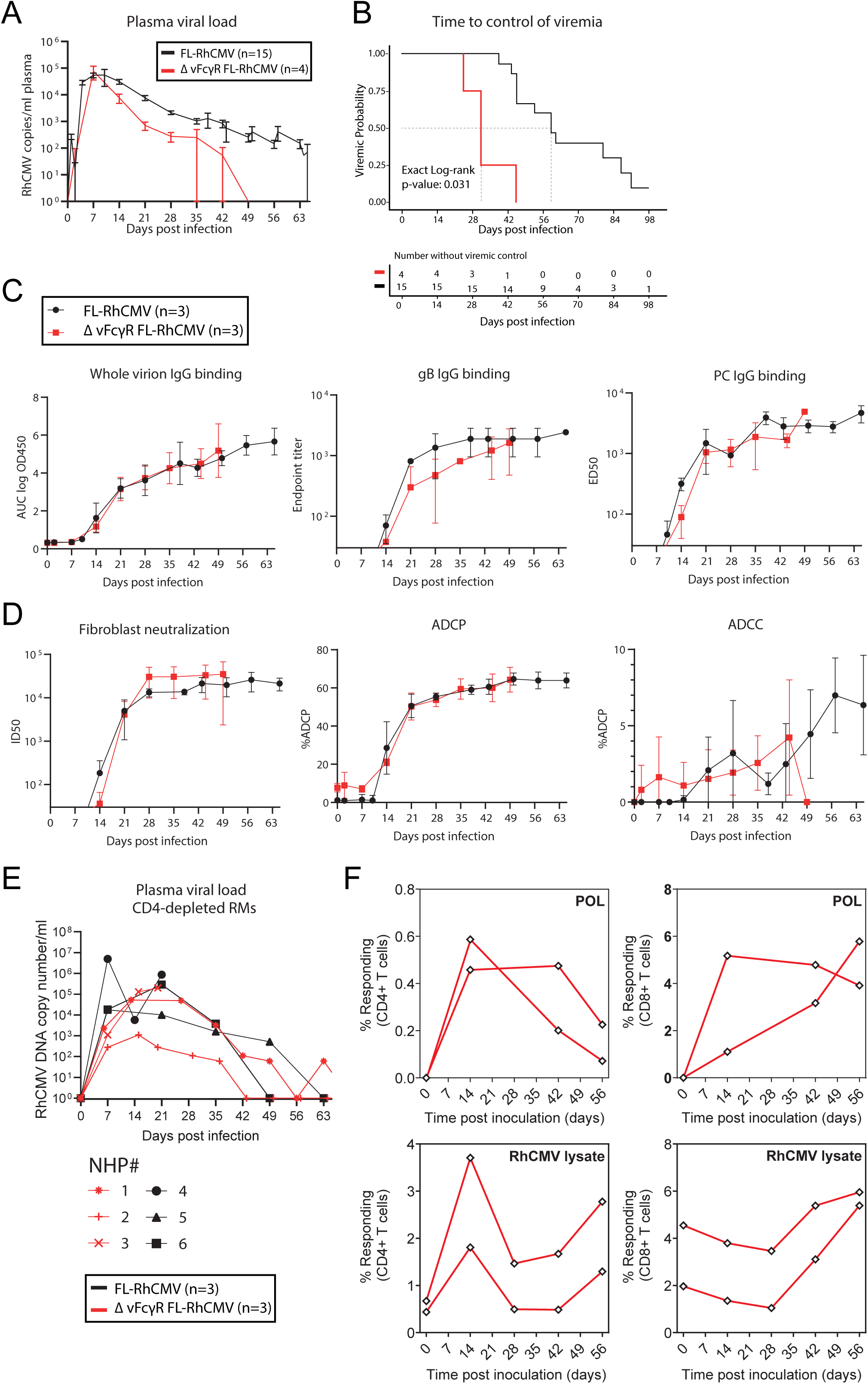
RhCMV encoded vFcγRs are essential for maintenance of viremia *in vivo*. A) Four RhCMV-seronegative male RM were inoculated with a FL-RhCMVΔΔΔ. As a control, three RhCMV-seronegative male RM and 12 RhCMV-seronegative female RM were inoculated with a FL-RhCMV. DNA was isolated form the plasma samples. Viral genome copy number were determined by qPCR using primer/probe sets targeting exon 1 of the IE locus. Copy numbers are reported via interpolation of a standard curve. B) Kaplan-Meyer survival analysis demonstrating the timing of control of viremia, defined as the first time point at which RhCMV is below the LOD (1 copy/well). The median time to control was 31.5 days post infection in animals infected with vFcγR-deleted RhCMV compared to 59.5 days in animals infected with FL-RhCMV (p = 0.031). C) IgG binding to whole FL-RhCMV virions, RhCMV gB, and RhCMV pentamer was measured by ELISA. D) Neutralization of FL-RhCMV on fibroblasts. ADCP is reported as the percentage of live THP-1 cells containing fluorescently labeled FL-RhCMV after incubation with plasma. ADCC is reported as the percentage of live rhesus CD16-expressing NK92 cells expressing the degranulation marker CD107a following co-incubation with FL-RhCMV infected fibroblasts and plasma samples. All responses demonstrate comparable magnitude and kinetics between animals infected with FL-RhCMV (n = 3) versus vFcγR-deleted RhCMVΔΔΔ (n = 4). E) Viral load kinetics in plasma of CD4+ T cell-depleted RhCMV-naïve animals infected with vFcγR-deleted RhCMV (black) versus FL-RhCMV (red). F) Two RhCMV-seropositive RMs were inoculated with 5×10^6^ PFU of vFcγR-deleted RhCMVΔΔΔ carrying an SIV-5’pol transgene as an immunological marker replacing the Rh13.1 ORF. The onset of SIV 5’Pol-specific CD4+ and CD8+ T cell responses and the boosting of RhCMV-specific T cell responses were measured in peripheral blood by intracellular cytokine staining (ICS) for IFN-γ and TNF-α using either a pool of 15mer peptides overlapping by 11 amino acids (AA) corresponding to SIV-5’pol (*52*) or RhCMV lysates. The frequency of IFN-γ+ and/or TNF-α+ memory T cells is shown for each individual animal and each indicated time point post-inoculation.

## Discussion

The goal of this study was to identify and characterize the full set of vFcγRs encoded by RhCMV in order to enable their assessment *in vivo* regarding interference with antibody function in a highly relevant animal model for HCMV. In addition to the previously identified Rh05, we now show that Rh152/151 and Rh173 also bind to host IgG and interfere with host FcγR activation by immune complexes. *Rh152/151* was identified by sequence homology to *UL119/118*. Remarkably, clear sequence orthologs of this gene can be found in all examined old and new world NHP CMV genomes and a non-spliced UL119/118-ortholog also displaying IgG-binding *in vitro* has even been identified in guinea pig CMV (GPCMV) (*27*), indicating that the ancestral gene predating all of these modern relatives was likely acquired by a common ancestor millions of years ago. Similar to HCMV UL119/118 (gp68), we observed that Rhesus Rh152/151 is able to directly interfere with the binding of host FcγRs to antibody complexes, indicating that it binds a similar region on IgG. In HCMV this interference does not seem to be a direct competition for IgG interaction for the same binding site since UL119/118 attaches to the CH2-CH3 interdomain region of Fcγ with high affinity whereas host FcγRs bind Fcγ in the CH2 domain and in the CH1-CH2 hinge implying an effect on Fcγ conformation and accessibility (*45*).

A systematic screen of all *RL11* family members encoded by RhCMV for IgG binding identified Rh173 as an additional vFcγR. We focused on this viral gene family as three of four HCMV vFcγRs, RL11 (gp34), RL12 (gp95) and RL13 (gpRL13), as well as the previously identified RhCMV encoded vFcγR Rh05 belong to this family. Intriguingly, both HCMV RL13 and its RhCMV ortholog Rh13.1 restrict virus replication *in vitro* and are hence selected against (*33*). However, only RL13 is also able to bind Fcγ *in vitro.* Within the *RL11* family, *Rh173* exhibits a high degree of sequence variability across RhCMV isolates consistent with continuous immune pressure, but despite this diversity, all tested Rh173 variants demonstrated equivalent vFcγR activity. This extraordinary level of sequence diversity is shared between Rh173 and its functional HCMV ortholog RL12 (gp95).

All identified vFcγRs are expressed with similar E/L kinetics during virus replication, but their subcellular localization in the infected cell varies significantly. While both Rh152/Rh151 and Rh05 demonstrate a punctate staining pattern in the infected cell implying their localization to sub-cellular vesicular compartments, they do not seem to co-localize and likely locate to the Golgi and lysosomes/autophagosomes, respectively. Yet, deletion of individual vFcγRs revealed that both Rh05 and Rh152/151 seem to interfere with host FcγR activation cooperatively, similar to previously reported results for their HCMV orthologs (*34*). This observation indicates that these protein might interact transiently e.g. on the infected cell or virion membrane. Deletion of Rh173 alone on the other hand, which largely co-localizes with Rh152/Rh152, only partially restored activation of the high affinity FcγR CD64. Further work will be required to determine the details of this potential cooperation.

Deletion of the full set of vFcγRs did not affect replication or tropism *in vitro*, yet it eliminated viral interference with host FcγR activation. These results indicate that RhCMV vFcγRs do not seem to play a significant role in viral entry, replication and release in tissue culture. This conclusion is also supported by the observation that peak viremia following infection of seronegative RM was not different between vFcγR-deleted FL-RhCMV and vFcγR-intact FL-RhCMV. Therefore, RhCMV vFcγRs do not seem to be required to overcome the immediate, innate host response to infection *in vivo*. This is in contrast to viral proteins such as Rh159 (UL148) that prevent the activation of NK cells through sequestration of NK cell activating ligands for NKG2D in virally infected cells (*46*). Interestingly, the murine CMV vFcγR m138 also downregulates NKG2D-ligands and, as a result, affects viral replication *in vivo*, even before the onset of adaptive immune responses (*47–51*). In contrast, no additional functions have so far been found for primate CMV vFcγRs.

It was somewhat unexpected that deletion of all vFcγRs from FL-RhCMV did not prevent superinfection of RhCMV seropositive RM as indicated by the development of *de novo* T cell responses to the inserted SIV antigen. These data are consistent with prior results showing that Rh05 is not required for reinfection (*28*). In contrast, viral evasion of CD8+ T cells seems to be essential to overcome pre-existing immunity (*52*). Thus it seems that evasion of humoral immunity may be a lesser requirement to overcome compared to T cell immune evasion of preexisting immune responses during superinfection.

The deletion of vFcγRs also did not affect the initial phases of primary infection. However, there was a significant impact on viral clearance from circulation that coincided with the development of a robust anti-viral antibody response. Notably, this analysis does have limitations in the small number of only male animals infected with vFcγR-deleted FL-RhCMV and the inclusion of both male and female animals infected with differing formulations of vFcγR-intact RhCMV. However, we observe remarkably consistent viral load kinetics among both vFcγR-intact RhCMV subgroups, supporting the validity of the observed difference in time to clearance. This difference between vFcγR-intact and -deleted FL-RhCMV was not observed upon deletion of CD4+ T cells from CMV naïve RM prior to inoculation. Since CD4+ T cells are required for the timely development of antibody responses and depletion of CD4+ T cells resulted in a delay of RhCMV-specific antibody responses (*12*), this result strongly suggests that a vFcγR-deleted RhCMV is more susceptible to antibody control than wildtype RhCMV, consistent with a role in evading the humoral response. In future experiments, we plan to compare infection of B-cell depleted animals which is expected to further support this immune evasion function.

Targeting vFcγRs therapeutically, e.g. by using antibodies that block their Fcγ-binding activities, could potentially increase the ability of anti-viral antibodies to control viral replication. Previously, we showed that pre-treatment of pregnant animals with HIG from RhCMV-positive animals prevented congenital transmission of RhCMV in immunosuppressed, RhCMV-seronegative animals (*12, 13*). However, this protective effect was only observed when the antibodies were isolated from donors with potent plasma virus neutralizing activity. The results presented here raise the intriguing possibility that vFcγRs play a role in limiting the activity of antibody therapies. Blocking vFcγR function, e.g. by antibody Fab fragments that prevent Fcγ-binding, could thus improve the ability of passively administered monoclonal or polyclonal antibodies to control viral replication in immunocompromised individuals, and possibly interrupt transmission to the fetus or ameliorate fetal disease. Further, inclusion of vFcγRs as antigens in an HCMV vaccine candidate may generate antibody responses that would block vFcγR-based inhibition of protective vaccine-elicited humoral immunity. Given our established RM model of congenital CMV transmission (*12*), we now have the unique opportunity to test these hypotheses in a relevant animal model for human vaccine development.

## Material and Methods

### Cells

All cells were cultured in a 5% CO_2_ atmosphere at 37°C. Telomerized rhesus fibroblasts (tRF), HEK293T cells, 293T-CD20 (kindly provided by Irvin Chen, UCLA (*53*)) and Hela cells were maintained in Dulbecco’s modified Eagle’s medium (DMEM, Gibco) supplemented with 10% (vol/vol) fetal calf serum (FCS, Biochrom) and antibiotics (1x Pen/Strep, Gibco). tRF were generated from primary rhesus fibroblasts (RF) obtained from animals housed at Oregon National Primate Research Center (ONPRC) and life-extended as described previously (*54*). BW5147 mouse thymoma cells (kindly provided by Ofer Mandelboim, Hadassah Hospital, Jerusalem, Israel) were maintained at 3×10^5^ to 9×10^5^ cells/ml in Roswell Park Memorial Institute medium (RPMI GlutaMAX, Gibco) supplemented with 10% (vol/vol) FCS, antibiotics, sodium pyruvate (1x, Gibco) and β-mercaptoethanol (0.1 mM, Gibco). Rhesus retinal pigment epithelial (RPE) cells were a kind gift from Dr. Thomas Shenk (Princeton University, USA) and were cultured in a mixture of DMEM and Ham’s F12 at 1:1 ratio supplemented with 5% FBS, antibiotics (1× Pen/Strep, Gibco), 1 mM sodium pyruvate, and nonessential amino acids.

### Cloning of RhCMV encoded vFcγRs and vFcγR candidates into expression plasmids

Sequences of RhCMV vFcγRs were synthesized as gBlock fragments (IDT) and cloned via Pst1 and BamH1 digest into a p-IRES-eGFP vector (Addgene) providing a unified tapasin signal peptide and resulting in an N-terminal HA-epitope tag upon cleavage of the tapsin signal peptide (*55*). Original signal peptides were predicted using SignalP 5.0 (DTU). CD4-tailed constructs in which the C-terminal TM and cytoplasmic domains were replaced with those of human CD4, were designed as described previously (*34*) and synthesized as gBlocks. A self-cleaving T2A peptide sequence was introduced into the 3’ end of the RhCMV ORF replacing the stop codon as described elsewhere (*37*). gBlock fragments were designed to encode Rhesus-CD4 (GenBank: M31134.1) and fused to the individual ORFs via the T2A peptide by insertion downstream of the ORFs cloned into p-IRES-eGFP.

### Immunoprecipitation of purified Fab, Fc and IgG from metabolically labeled cells

tRFs were seeded at 1×10^6^ cells per well (6-well plate) and infected at a multiplicity of infection (MOI) of 5 with the RhCMV strains 68-1, 68-1R or mutants thereof. Cells were labelled with [^35^S]-Met/Cys for 2 h at 72 hpi. After NP-40 cell lysis, immunoprecipitation of proteins was performed by incubation of lysates with 10 µg/ml anti-Rhesus-CD4 IgG1 (Nonhuman Primate Reagent Resource Cat#PR-0407) for 1h followed by retrieval of proteins using protein A Sepharose beads (PAS). Proteins were selectively deglycosylated using Endo H or PNGasF digestion and subsequently separated on a 10% - 13% SDS-PAGE gradient. Metabolic labelling was visualized using a Typhoon phosphoimager (Cytiva).

### Parental virus strains and construction of recombinants

All RhCMV recombinants generated for this study were based on bacterial artificial chromosomes (BAC) of either the tissue culture adapted RhCMV 68-1 strain (GenBank #MT157325) or full-length (FL)-RhCMV (GenBank #MT157327) which was constructed by reversing genomic inversions, deletions and ORF truncations identified in 68-1 (*2*). The FL-RhCMV clone now contains a complete genome representative of primary, low passage viral isolates and demonstrates enhanced virulence and replication in immunocompetent RM when compared to 68-1 (*2*). FL-RhCMV-based recombinants either contained riboswitches introduced into the 5’ and the 3’ flanking regions the Rh13.1 ORF (*2*), or replacements of the entire Rh13.1 ORF with heterologous antigens derived from simian immunodeficiency virus (SIV). Since intact Rh13.1 impairs growth in vitro, similar to HCMV RL13 (*56, 57*), these modifications allow for high-titer stock production without selection against the Rh13.1 ORF (*2*). Additionally, we used the inserted antigens as immunological makers after inoculating CMV-positive animals since we can monitor the development of antigen-specific T cell responses as an indicator for infection and re-infection.

FL-RhCMVΔRh13.1/SIVgag/Rh05-HA/Rh173-V5, FL-RhCMVΔRh13.1/SIVgag/ΔRh05/ΔRh152/152ΔRh173, and FL-RhCMVΔRh13.1/SIV5’pol/ΔRh05/ΔRh152/152ΔRh173 were all based on RhCMVΔRh13.1/SIVgag, which has been described before (*41*). HA and V5 were added to Rh05 and Rh173 through sequential homologous *en passant* recombination using specific primers pairs containing the DNA sequence for the peptide tags. The triple vFcγR deletion mutant (ΔΔΔ) was based on FL-RhCMVΔRh13.1/SIVgag and constructed by sequentially deleting the entire coding region of each ORF. For superinfection experiments using the ΔΔΔ recombinant in RhCMV seropositive RM, SIVgag was replaced with SIV5’pol. RhCMV 68-1R was constructed by repairing the SNP resulting in a premature termination codon in Rh152/Rh151 by homologous *en passant* recombination. The same techniques was then used to either create single deletion mutants of individual vFcγR (Rh05, Rh152/151 or Rh173) or a triple (ΔΔΔ) vFcγR deletion mutants in which all ORFs were sequentially fully deleted.

The *en passant* recombination technique (*58*) used to introduce alterations into the parental BACs had been previously adapted to our RhCMV vector system (*41*). In short, this technique requires the duplication of a 50-bp stretch of the genome flanking the targeted genomic locus. This duplication can be achieved through strategic primer design. These primers are then used to amplify an aminoglycoside 3-phosphotransferase gene conferring kanamycin resistance (KanR) as a selection marker preceded by an I-SceI homing enzyme target sequence. After homologous recombination, the selection marker is flanked by the introduced direct repeats. Expression of I-Sce I by arabinose induction in E.coli strain GS1783 harboring the RhCMV BACs results in the selective induction of DNA double-strand breaks in the RhCMV BAC. These breaks are then repaired via recombination of the repeated DNA sequences by inducing the expression of the λ phage-derived Red recombination genes through heat shock. This results in complete removal of the KanR resistance cassette without retaining any DNA sequences introduced during recombineering. Final constructs were analyzed by Xma I restriction digest, Sanger sequencing of the altered genome region, and full-genome analysis by next generation sequencing (NGS) before virus reconstitution.

### Multistep Growth Curves

Primary rhesus fibroblasts (RF) were seeded at 5 x 10^4^ cells per well in 24-well plates (Corning). On the next day, the cells were infected in triplicate with either FL-RhCMV ΔRh13.1/SIVgag or the FL-RhCMV ΔRh13.1/SIVgag ΔRh05ΔRh152/Rh151ΔRh173 (ΔΔΔ) recombinant at a multiplicity of infection (MOI) of 0.01. Starting from day 3 post infection, supernatants were collected every third day until day 24 and stored at -80°C. To determine the number of focus forming units (FFU) in each collected sample, 10^4^ RF per well were seeded into 96-well plates (Corning) in quadruplicates and infected the next day with the supernatant samples. At 72 hpi, the cells were fixed with 100% methanol for 20 minutes at - 20°C and then permeabilized with 2% paraformaldehyde and 0.1% Triton X-100 in Phosphate Buffered Saline (PBS) for 15 min at room temperature (RT). Subsequently, the cells were blocked for 30 min with 2% BSA in PBS (PBA) and stained with a mouse anti-RhCMV pp65b mAb (clone 19C12.2, generated in house by OHSU monoclonal antibody shared resource (*59*)) for 1 h at 37°C. After washing three times with PBA, the cells were stained with an Alexa488-conjugated anti-mouse secondary antibody (Invitrogen) for 1h at 37°C and then with 3 µM DAPI (Invitrogen) for 20 min. Images were acquired using an EVOS fluorescence microscope (Life Technologies) and the ImageJ software was used to process the images and count all pp65b+ and DAPI+ cells. Double positive versus single positive (DAPI+) cell counts were used to calculate the virus titer for each sample as focus forming units per milliliter (FFU/ml).

### Quantitative reverse transcriptase PCR (qRT-PCR) analysis to determine the kinetics of RhCMV Fc**γ**Rs transcripts

RF were seeded in 6-well plates and infected with either FL-RhCMV or FL-RhCMV ΔRh05ΔRh152/151ΔRh173 (both also containing ΔRh13.1/SIVgag) as a negative control at a MOI of 5. Subsequently, total RNA was isolated using TRIzol reagent (Thermo Fischer Scientific) at 4, 8, 12, 24, 36, and 48 h post infection (hpi) following the manufacturer’s instructions. Next, 1 μg of total RNA per sample was DNase I (Zymo Research) treated and then used to synthesize complementary DNA (cDNA) using the Maxima Reverse Transcriptase (Thermo Fisher Scientific). To determine the transcription levels and temporal kinetics of the RhCMV FcγRs *Rh05, Rh152/151, and Rh173*, qPCR was performed with the cDNA using TaqMan Fast Advanced Master Mix (Applied Biosystems) and QuantStudio 7 Flex Real-Time PCR Systems (Applied Biosystems). Transcript copy numbers of target genes were determined using the QuantStudio Real-Time PCR Software v1.3 and then normalized to the housekeeping transcript for Glyceraldehyde 3-phosphate dehydrogenase (GAPDH). Data were graphed for each gene and time point as relative mRNA copy numbers. To determine the kinetic class of each vFcγR transcript, the immediate early (IE) gene *Rh156*, the early (E) gene *Rh189,* and the late (L) gene *Rh137* were included as controls as described previously (*41*). For the q-PCR assay, primers and probes specific to each gene of interest were used (Table S1). Expression of genes adjacent to the deleted genes in FL-RhCMV ΔRh05ΔRh152/151ΔRh173 was determined by qualitative PCR using the same cDNA and primer sets specific to each deleted gene and the corresponding adjacent genes (Table S1). To confirm the absence of residual viral DNA in the cDNA samples, a set of RNA samples was used during cDNA synthesis without reverse transcriptase (RT-) treatment.

### RhCMV virion isolation and virion proteins detection by immunoblot

FL-RhCMVΔRh13.1/SIVgag, FL-RhCMVΔRh13.1/SIVgag/ΔRh05/ΔRh152/152ΔRh173 (ΔΔΔ) and FL-RhCMVΔRh13.1/SIVgag/Rh05-HA/Rh173-V5 virions were purified over a discontinuous Nycodenz gradient, as described previously (*59*). The virus was isolated from the culture medium of infected RF. For each virus, 10 T-175 flasks of confluent RFs were infected at a MOI of 0.05-0.1. Infected cells were harvested after all cells showed maximal cytopathic effect. The supernatants were first clarified by centrifugation at 7,500 g for 15 min. The clarified medium was layered over a sorbitol cushion (20% D-sorbitol, 50 mM Tris [pH 7.4], 1 mM MgCl_2_), and virus was pelleted by centrifugation at 64,000 g for 1 h at 4°C in a Beckman SW28 rotor. The virus pellet was resuspended in TNE buffer (50 mM Tris [pH 7.4], 100 mM NaCl, and 10 mM EDTA). The virus particles were further purified by layering them over a discontinuous 5% to 50% Nycodenz gradient (Sigma-Aldrich) in TNE buffer, followed by centrifugation at 111,000 g for 2 h at 4°C in a Beckman SW41 Ti rotor. The virion bands in the gradient were extracted with a syringe through the side of the centrifuge tube, and the particles were pelleted in a Beckman TLA-45 rotor in a Beckman Optima TL 100 Ultracentrifuge at 40,000 g for 1 h and washed twice with TNE buffer. In order to detect RhCMV FcγRs in the RhCMV virion preparation, pellets were lysed with 200 µl RIPA lysis buffer (ThermoFischer Scientific) and 200 µl of 2x Laemmli buffer (0.125M Tris HCl pH6.8, 4% SDS, 20% glycerol, 0.004% bromophenol blue). Protein lysates were denatured at 95°C for 10 min and similar protein amounts were electrophoretically separated on a NuPAGE™ 10%, Bis-Tris gel (Invitrogen, ThermoFischer Scientific), electrophoretically transferred onto PVDF transfer membranes (ThermoFischer Scientific) and stained for HA (Sigma-Aldrich, HA-7), V5 (Invitrogen, E10/V4RR), Rh152/151 , and Rhpp65b using specific antibodies. Rh152/151 and Rhpp65b were made by OHSU monoclonal antibody shared resource (*42*). Protein bands were visualized by SuperSignal^TM^ West pico PLUS chemiluminescent substrate (ThermoFischer Scientific).

### Comparison of fibroblasts and epithelial cell infections

RF and rhesus retinal pigment epithelial cells (RPE) were seeded at 12,000 cells per well onto 96-well plates. After 24 h, the cells were infected with either FL-RhCMV, vFcγRs deleted FL-RhCMV (ΔΔΔ), or pentameric complex (PC)-deleted FL-RhCMV (deleted for Rh157.5 and Rh157.4, the homologues of HCMV UL128 and UL130, all ΔRh13.1/SIVgag) at a 1:3 serial dilution starting at 1:50 inoculum in DMEM complete. On day 72 hpi, all cells were fixed with methanol for 20 min at -20°C, then washed three times with PBS. Subsequently, fixed cells were stained with the mouse α-RhCMV pp65b antibody (clone 19C12.2, generated in house by OHSU monoclonal antibody shared resource) for 1 h at 37°C. Next, cells were stained with Alexa488-conjugated anti-mouse secondary antibody (Thermo Fisher) for 1 h at 37°C and then with DAPI (Thermo Fisher) for 10 min at RT. Images were acquired using the EVOS® FL Auto Imaging System (Thermo Fisher) and analyzed using the ImageJ software. Relative infection rates were calculated as the ratio of focus forming units per ml (FFU/ml) in RPE cells versus FFU/ml in RF for each recombinant. Experiments were performed in triplicate in three biological repeats.

### Immunofluorescence assay and fluorescence microscopy

tRFs were cultured in DMEM supplemented with 10% fetal bovine serum (FBS), 100 U/ml penicillin, and 100 µg/ml streptomycin. 10^6^ TRFs were plated overnight on 12 mm wide, 0.13-0.17 mm thick glass coverslips in 24-well plates. The next day cells were infected with FL-RhCMV, FL-RhCMVΔRh152/152 or FL-RhCMV/Rh05-HA/Rh173-V5 at MOI of 1. At 24 hpi and 48 hpi, cells were rinsed with PBS and fixed with 4% methanol-free formaldehyde for 10 min at RT. Cells were washed thrice using PBS and blocked and permeabilized using saponin-BSA buffer for 1 h at RT (0.2% saponin from Millipore and 1% BSA from Fisher). Cells were incubated with the following primary antibodies for 1 h at RT: Anti-HA (Abcam-Ab18181), Anti-RhCMV-Rh152/151 (mouse mAb generated in house by OHSU monoclonal antibody shared resource (*42*)). Anti-V5 (GeneTex-GTX628529), Anti-RhCMV IE2 (mouse mAb generated in house by OHSU monoclonal antibody shared resource (*42*)), Anti-EEA1 (CST-2411S), Anti-LAMP1 (Thermo-Fisher, PA1-654A), Anti-Rab6A (Thermo-Fisher, PA5-22127), Anti-ERp57 (Invitrogen-PA3-009). Coverslips were washed thrice for a total of 30 min using saponin-BSA buffer. Secondary antibodies (A21463 anti-mouse 647, A21441 anti-rabbit 488, A21155 IgG3-594, A21131 IgG2a 488, A21240 IgG1 647, all from Thermo-Fisher) and DAPI (D1306, Thermo-Fisher) in saponin-BSA buffer were incubated for 1 h at RT. Cells were washed thrice for a total of 30 min in saponin-BSA buffer followed by a final wash using PBS. Coverslips were mounted on glass slide using Prolong Gold (P36930, Thermo-Fisher) and cured for 24-48 h. The cells were imaged using the 100x oil immersion objective of a Keyence BZ-X710 microscope using its built-in 2.8 megapixel monochrome CCD camera. To eliminate fluorescence blurring in images, sectional function of 2D structured illumination and haze reduction features were used. Data was processed using FIJI software. All the immunofluorescence experiments were independently repeated four times.

### Flow cytometry

Cells were washed in PBS, equilibrated in staining buffer (PBS, 3% FCS) and pelleted at 1000 g and 10°C for 3 min. Cells were incubated in staining solution (antibody diluted in buffer as suggested by the supplier). Every incubation step was carried out at 4°C for 1h, followed by 3 washing steps in staining buffer. Dead cells were stained using DAPI. Cells were analyzed on a FACS Fortessa LSR instrument (BD Bioscience). Intracellular stains were performed using the Cytofix/Cytoperm kit according to the supplier’s instructions (Becton Dickinson). HA-epitopes were detected using PE-labeled clone 16B12 (Biolegend). PE-conjugation was performed using an ab102918 labelling kit (Abcam) as suggested by the supplier. Data was evaluated using FlowJo 10 software.

### Fc**γ**-receptor activation assay

The assay was performed as described previously (*28, 34, 39*). Briefly, target cells were incubated with dilutions of a rhesus-CD4 specific rhesusized IgG1 (*28*) or a serum pool from 8 RhCMV-seropositive rhesus macaques in DMEM supplemented with 10% (vol/vol) FCS for 30 min at 37°C. Cells were washed before co-cultivation with FcγR-activation reporter cells (ratio E:T 10:1) for 16 h at 37°C in a 5% CO_2_ atmosphere. For all activation assays, mouse IL-2 secretion was quantified by anti-IL-2 ELISA as described previously (*39*).

### CD16 binding assay

The assay was performed as described previously (*34*). In brief, 293T-CD20 cells expressing transfected vFcγRs or control constructs were harvested using Accutase® (Sigma-Aldrich) to retain surface molecules upon detachment. Harvested cells were washed in PBS, equilibrated in staining buffer (PBS, 3% FCS) and sedimented at 1000g and 10°C for 3 min. Cells were then incubated with staining buffer containing a humanized human CD20-specific mAb (rituximab) followed by incubation with recombinant human or rhesus FcγR ectodomains pre-incubated with a His-tag-specific PE-labeled mAb. His-tagged FcγR ectodomains were used at a final concentration of 5 µg/ml (Sino Biological). Dead cells were labeled via DAPI stain. Analysis was performed on a FACS Fortessa LSR instrument (BD Bioscience).

### Rhesus macaques

Indian-origin rhesus macaques were housed at the Tulane National Primate Research Center, the California National Primate Research Center or the Oregon National Primate Research Center and maintained in accordance with institutional and federal guidelines for the care and use of laboratory animals (National Research Council Guide for the Care and Use of Laboratory Animals. National Academies Press, 8th Ed, Washington, DC, 2011). RhCMV-seronegative male RM were infected with 1×10^6^ plaque forming units (pfu) of either FL-RhCMV (n = 3) or vFcγR-deleted FL-RhCMVΔΔΔ (n = 4). Blood samples were collected at 0, 2, 7, 10, 14, 21, 28, 38, 43, 50, 57, and 64 or 65 days post infection (dpi) for FL-RhCMV infected animals and 0, 2, and 7 dpi and weekly thereafter through 6-7 weeks post infection for ΔvFcγR RhCMV infected animals. Blood samples were collected at weeks 0, 2, 6, 8. An additional 12 pregnant RhCMV-seronegative RM infected with 1×10^6^ PFU each of FL-RhCMV and UCD52 in late first/early second trimester were included as additional controls to show viral kinetics in animals infected with vFcγR-intact viruses. The dams were sampled at days 0, 1, 4 post infection and then on a weekly basis thereafter. To deplete CD4+ T cells, six dams received an 50 mg/kg intravenous dose of recombinant rhesus anti-CD4 antibody (CD4R1 clone; Nonhuman Primate Reagent Resource) one week prior to infection with 1×10^6^ pfu of either FL-RhCMV (n = 3) or FL-RhCMVΔΔΔ (n = 3) in late first/early second trimester. Blood was collected on an approximately weekly basis starting at anti-CD4 infusion through 12 weeks post infection for these dams. For re-infection studies, an additional two seropositive rhesus macaques we inoculated with 1×10^7pfu vFcγR-deleted FL-RhCMVΔΔΔ.

### Viral load qPCR

DNA was isolated using the QIAmp DNA minikit (Qiagen). We utilized a 40-cycle real-time qPCR reaction using 5′-GTTTAGGGAACCGCCATTCTG-3′ forward primer, 5′-GTATCCGCGTTCCAATGCA-3′ reverse primer, and 5′-FAM-TCCAGCCTCCATAGCCGGGAAGG-TAMRA-3′ probe directed against a 108bp region of the highly conserved RhCMV *IE* gene, using a standard IE plasmid to interpolate the number of viral DNA copies/ml of plasma. Two out of six replicates above the limit of detection (1-10 copies per well) was required to report a positive result for each plasma sample.

### Antibody serology assays

RhCMV-specific IgG antibody kinetics were measured in plasma by enzyme-linked immunosorbent assays (ELISAs) using either whole virion preparations of FL-RhCMV or purified glycoprotein preparations of gB or pentamer as described previously (*13*). Briefly, high-binding 384-well ELISA plates (Corning) were coated with 15 μl/well of 5120 pfu/ml RhCMV or 2μg/ml purified gB or pentameric complex (PC) protein in 0.1M sodium bicarbonate (pH = 9.55) overnight at 4°C. Following overnight incubation, plates were blocked for 1-2 h with blocking solution (PBS+, 4% whey protein, 15% goat serum, 0.5% Tween-20), and then 3-fold serial dilutions of plasma (1:30 to 1:65,610) were added to the wells in duplicate for 1-2 h. Plates were then washed using an automated plate washer (BioTek) and incubated for 1 h with mouse anti-rhesus IgG HRP-conjugated secondary antibody (Southern Biotech, clone SB108a) at a 1:5,000 dilution or anti-rhesus IgM HRP-conjugated secondary antibody (Rockland) at 1:8,000. After two washes, SureBlue Reserve TMB Microwell Peroxidase Substrate (KPL) was added to the wells for 7 min for whole virion ELISAs and 3.5 min for glycoprotein ELISAs. The reaction was stopped by addition of 1% HCl solution (KPL) and plates were read at 450 nm. The cutoff for positive antibody reactivity was considered to be 3 SDs above the average OD_450_ measured for RhCMV-seronegative samples at the starting plasma dilution (1:30). ED_50_ was calculated as the sample dilution where 50% binding occurs by interpolation of the sigmoidal binding curve using four-parameter logistic regression, and this value was set to half of the starting dilution (15) when the 1:30 dilution point was negative by our cutoff and when the calculated ED_50_ was below 15. Endpoint titer or area-under-the-curve (AUC) were used to report binding results when the dilution series resulted in incomplete sigmoidal curves, and thus unreliable ED_50_ calculations, for the majority of positive samples. Endpoint titer is defined as the last dilution factor at which there was binding above the positivity cutoff. AUC was calculated by integrating the OD_450_ over the full dilution series.

Neutralization of RhCMV on fibroblasts was monitored as previously described (*13*). Briefly, serial dilutions (1:30 to 1:65,610) of heat-inactivated rhesus plasma were incubated with FL-RhCMV (*2*) (MOI = 1) for 1–2 h at 37°C. The virus/plasma dilutions were then added in duplicate to 384-well plates containing confluent cultures of TRF cells and incubated at 37°C for 24 h. Infected cells were then fixed for 20 min in 10% formalin and processed for immunofluorescence with 1 μg/ml mouse anti-RhCMV pp65B or 5 ug/ml mouse anti-Rh152/151 monoclonal antibody followed by a 1:500 dilution of goat anti-mouse IgG-Alexa Fluor 488 antibody (Invitrogen). Nuclei were stained with DAPI for 10 min (Invitrogen). Infection was quantified in each well by automated cell counting software using a Molecular Devices ImageXpress Pico. Subsequently, the ID_50_ was calculated as the sample dilution that caused a 50% reduction in the number of infected cells compared with wells treated with virus only. If the ID_50_ was below the limit of detection (e.g., every value in the series resulted in a level of infection above 50%), we set the ID_50_ to 15 (half of the first dilution).

Antibody-dependent cellular phagocytosis was assessed by conjugation of concentrated FL-RhCMV to Alexa Fluor 647 using NHS-ester reaction (Invitrogen), which was allowed to proceed in the dark with constant agitation for 1–2 h and then quenched by the addition of pH 8.0 Tris hydrochloric acid, and 5,000 pfu of the conjugated virus (10 μl) of virus and plasma samples in duplicate at a 1:30 dilution were combined in a 96-well U-bottom plate (Corning) and incubated at 37°C for 2 h. THP-1 monocytes were then added at 50,000 cells per well. Plates were spun for 1 h at 1200xg at 4°C and then transferred to a 37°C incubator for an additional 1 h. Cells were then washed with 1% FBS in PBS and stained with aqua live/dead (Invitrogen) at 1:1000 for 20 min. Following another wash step, cells were fixed for 20 min in 10% formalin and resuspended in PBS. Fluorescence was measured using a BD Fortessa flow cytometer using the high-throughput sampler (HTS) attachment, and data analysis was performed using FlowJo software (v10.8.1). Results are reported as the percent of the live population that was AF647+.

Antibody-dependent cellular cytotoxicity was measured by NK cell degranulation using NK92.rh.158I.Bb11, an engineered NK cell line expressing the Ile158 variant of rhesus macaque CD16 (*60*). Confluent TRF cells were infected with 1 MOI FL-RhCMV in low serum (5%) medium in T75 flasks (ThermoFisher). Mock infection was performed in parallel. After 24 h of infection, cells were dissociated from the flask using TrypLE (Gibco), and 50,000 target cells were seeded in each well of a 96-well flat bottom tissue culture plate (Corning). Cells were incubated for 16–20 h to allow them to adhere. Then, the target cell medium was removed, and 50,000 NK92.Rh.CD16-Bb11 cells were added to each well with Brefeldin A (GolgiPlug, 1:1000, BD Biosciences), monensin (GolgiStop, 1:1500, BD Biosciences), CD107a-FITC (BD Biosciences, 1:40, clone H4A3), and plasma samples at 1:25 dilution or purified IgG controls from either seropositive or seronegative plasma donors at 50–250 μg/ml in RPMI1640 containing 10% FBS, plated in duplicate, and incubated for 6 h at 37°C and 5% CO_2_. The same samples were added to mock-infected target cells for background subtraction. NK cells were then collected and transferred to a 96-well V-bottom plate (Corning). Cells were pelleted and resuspended in aqua live/dead diluted 1:1000 for a 20 min incubation at RT. Cells were washed with PBS + 1% FBS and stained with CD56-PECy7 (BD Biosciences, clone NCAM16.2) and CD16 PacBlue (BD Biosciences, clone 3G8) for a 20 min incubation at RT. Cells were washed twice then resuspended in 10% formalin for 20 min. Cells were then resuspended in PBS for acquisition. Events were acquired on a BD Symphony A5 using the HTS attachment, and data analysis was performed using FlowJo software (v10.8.1). Data is reported as the % of CD107a+ live NK cells (singlets, lymphocytes, aqua blue–, CD56+, CD107a+) for each sample, background subtracted for first the no antibody control wells within mock and infected conditions, followed by subtracting signal from mock infected cells treated with the same sample.

### T cell assays

CD4+ and CD8+ T cell responses were measured in PBMC by flow cytometric intracellular cytokine staining (ICS) as described in detail previously (*41*). Frozen PBMCs were rapidly thawed at 37°C and washed with R10-media (RPMI 1640 medium (Thermo Fisher Scientific) supplemented with 10% Newborn Calf Serum (Gibco), 2mM l-glutamine (Thermo Fisher Scientific), 100U/ml penicillin (Invitrogen), 100µg/ml streptomycin (Invitrogen), 50µM βME (Thermo Fisher Scientific) before cells were combined with anti-CD28 (CD28.2, Purified 500ng/test: eBioscience), anti-CD49d mAb (9F10, Purified 500 ng/test: eBioscience), and test antigen then incubated for 1 h at 37°C before the addition of Brefeldin A, followed by an additional 8 h incubation. Co-stimulation without antigen served as a negative control. Cells were then stained with the following fluorochrome conjugated antibodies: anti-CD3 (SP34–2, Pacific Blue; BD Biosciences), anti-CD8 (SK1, PerCP-eFluor710; Life Tech), anti-CD4 (L200, BV510; BD Biosciences), anti-CD69 (FN50, PE-TexasRed, BD Biosciences), anti-IFNγ (B27, APC; BD Biosciences), anti-TNFα (MAb11, PE; BD Biosciences), and Ki67 (B56; FITC, BD Biosciences). T cell responses to SIV 5’Pol were measured by ICS using a mix of sequential 15-mer peptides (11 amino acid overlap). Stained samples were analyzed on an LSR-II or FACSymphony A5 flow cytometer (BD Biosciences) and analyzed using FlowJo 10 software (BD Biosciences). In all analyses, gating on the lymphocyte population was followed by the separation of the CD3+ T cell subset and progressive gating on CD4+ and CD8+ T cell subsets. Antigen-responding cells in both CD4+ and CD8+ T cell populations were determined by their intracellular expression of CD69 and either or both of the cytokines IFN-γ and TNF-α. Assay limit of detection was determined with 0.05% after background subtraction being the minimum threshold used in this study. After background subtraction, the raw response frequencies above the assay limit of detection were “memory-corrected” (i.e. percent responding out of the memory population), as previously described (*41*).

### Statistical analysis

Survival analysis comparing time to viremia stabilization, defined as the first timepoint post-peak viremia that plasma viral load (VL) was below the qPCR limit of detection, between rhesus macaques infected with FL-RhCMV and ΔvFcγR RhCMV was assessed via an exact log rank test using the coin package (*61, 62*). Kaplan-Meier estimates of viremic probabilities were employed for visualization and quantifying median time to viremia stabilization (*63*). All statistical analyses were performed using R statistical software (www.r-project.org).

## Acknowledgments

The authors would first like to acknowledge the rhesus macaques that were used in this study and thank the faculty and staff of the Departments of Veterinary Medicine and Collaborative Research at the Tulane National Primate Research Center (TNPRC) and the Oregon National Primate Research Center (ONPRC) for their excellent care of our research animals. We acknowledge the Molecular Virology Core at Oregon National Primate Research Center (ONPRC) for performing viral productions and qPCRs. We are additionally grateful to the NIH Nonhuman Primate Reagent Resource (R24 OD010976 and NIAID contract HHSN 272201300031C) which provided the CD4-depleting antibody. We further wish to acknowledge support from the Biostatistics, Epidemiology and Research Design (BERD) Methods Core. Fig. 1A and Fig. 4A cartoons were generated with Biorender (biorender.com), License University Medical Center Freiburg, Institute of Virology. We extend acknowledgements to Estes lab at VGTI for offering their Keyence microscope for generation of the IFA data.

## Author contributions

Conceived and designed the experiments: PK, DM, SRP, AK, KF.

Performed the experiments: CO, ME, SP, HT, KH, PK, MJM, EAS, LMS, SK, TDM, NHVB, MD, ZS.

Analyzed the data: PK, DM, CO, AD, RB, DS, SGH.

Writing and original draft preparation: PK, DM, CO.

Review and editing: SP, KF, HH, CC.

---

**Table S1.**
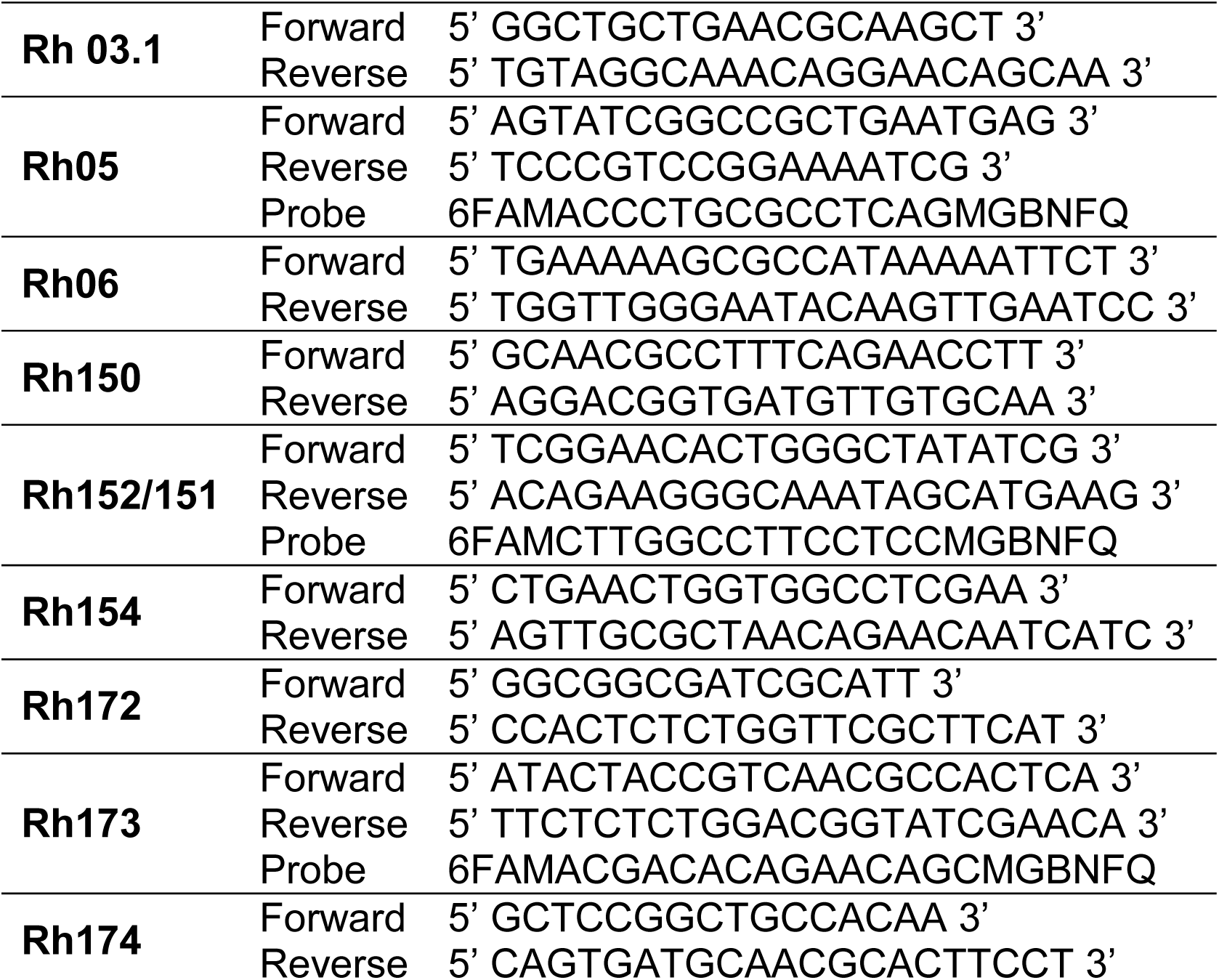
Primers and probes for the RT-qPCR/RT-PCR analyses.

**Fig S1.**
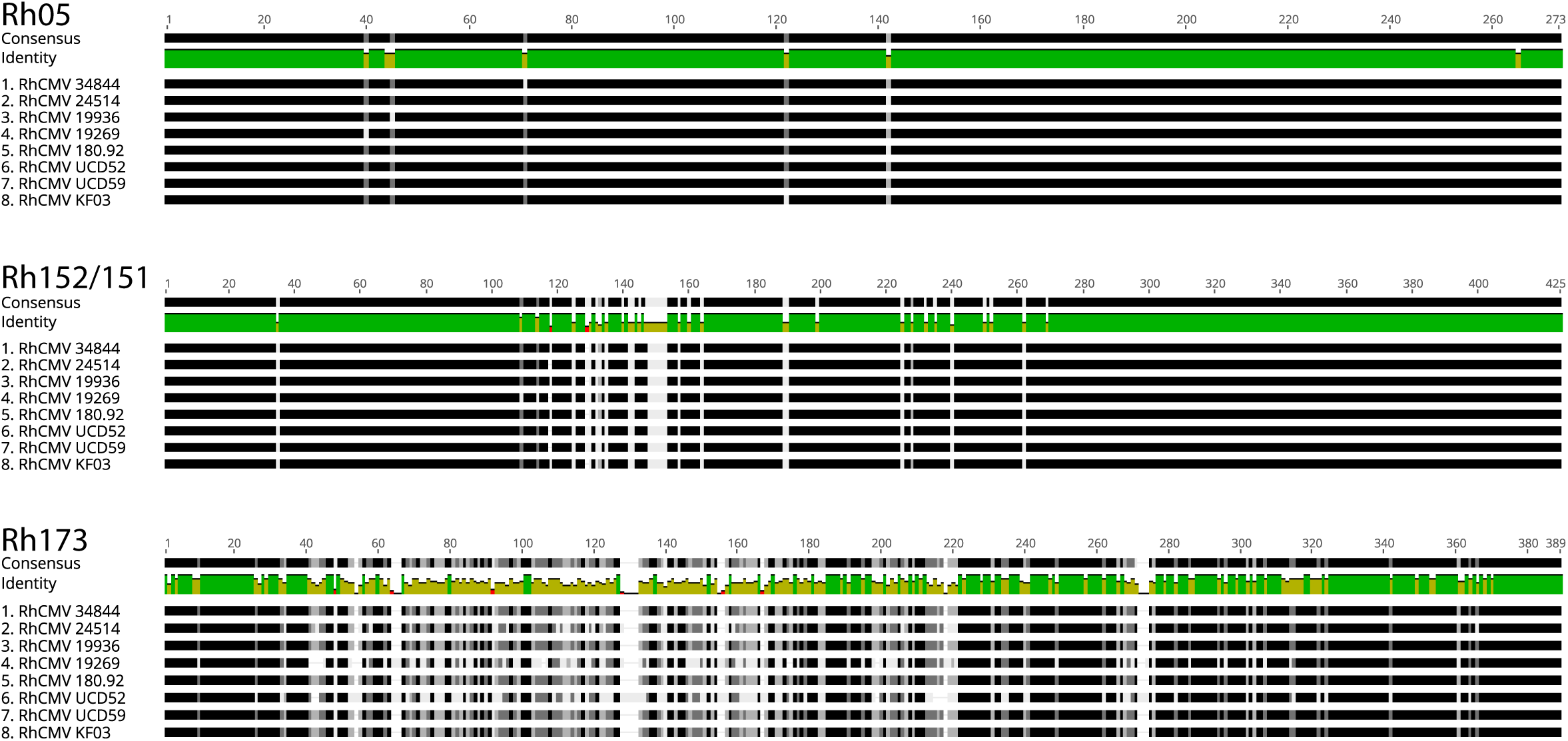
RhCMV vFcγR amino acid sequence alignments. Amino acid sequence alignments of RhCMV vFcγRs across all published full-length RhCMV genome sequences from primary isolates and excluding laboratory adapted strain sequences.

**Fig S2.**
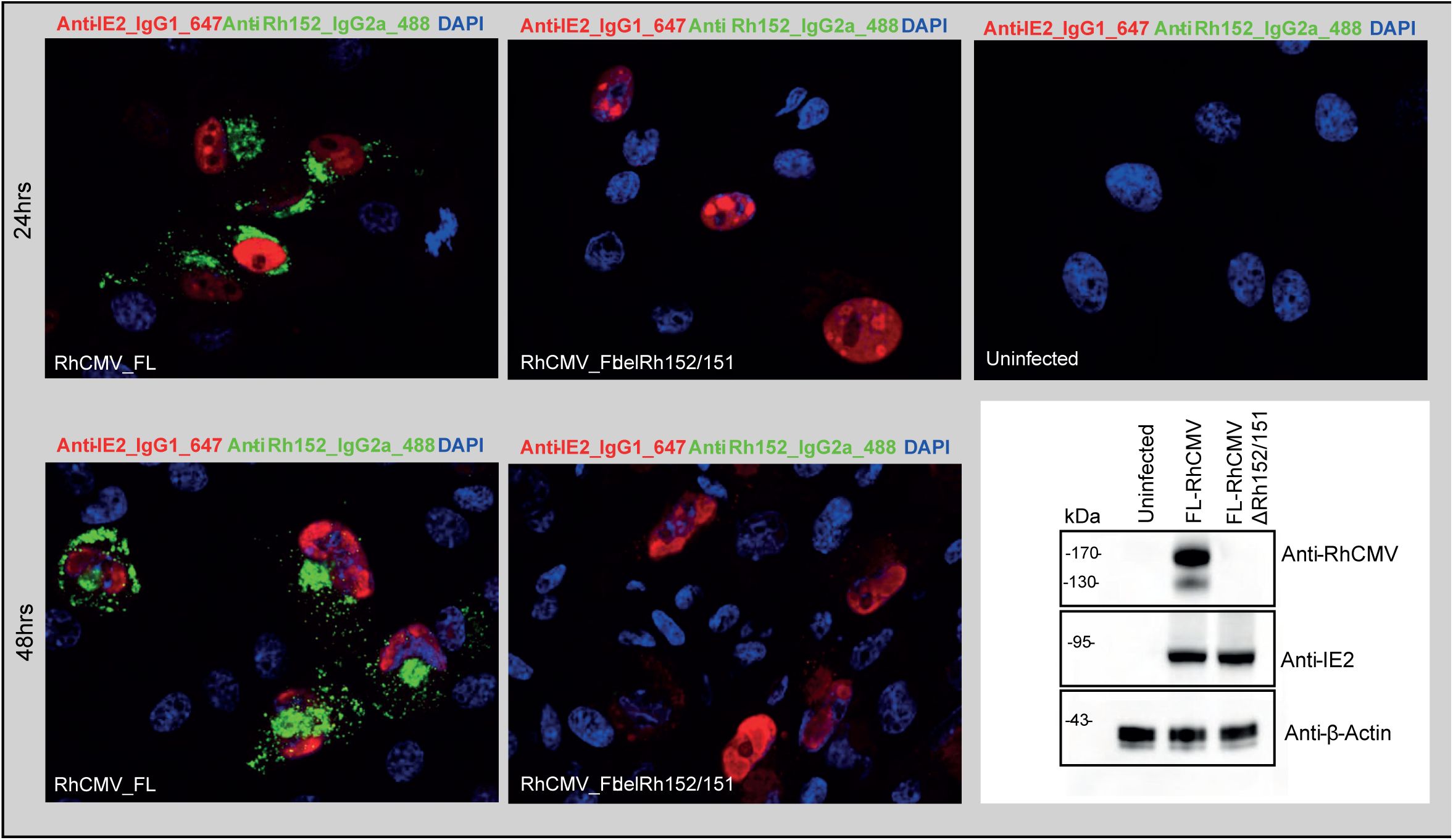
A RhCMV-specific mAb recognizes Rh152/151. A mAb from a screen of RhCMV specific mouse hybridoma cell lines (*42*) was tested for RhCMV protein specificity. tRFs were infected with FL-RhCMV or FL-RhCMVΔRh152/151 at MOI 1. IFA was performed at 24 hpi and 48 hpi using anti-IE2 and anti-RhCMV antibodies for detecting infected cells. Uninfected tRFs show absence of fluorescence signals, while at 24 hpi and 48 hpi cells show nuclear IE2 staining. The RhCMV-specific antibody displayed a punctate patterns at both time points. In contrast, FL-RhCMVΔRh152/151-infected cells displayed a loss of RhCMV-specific antibody binding, while preserving IE2 detection. This indicated that the RhCMV-specific antibody recognizes the Rh152/151 protein in RhCMV infected cells. This conclusion was confirmed by immunoblot analysis where the same RhCMV-specific mAb detected a protein band that was absent from FL-RhCMVΔRh152/151-infected cell lysates. IE2 and β-actin were included as infection and loading control, respectively.

**Fig S3.**
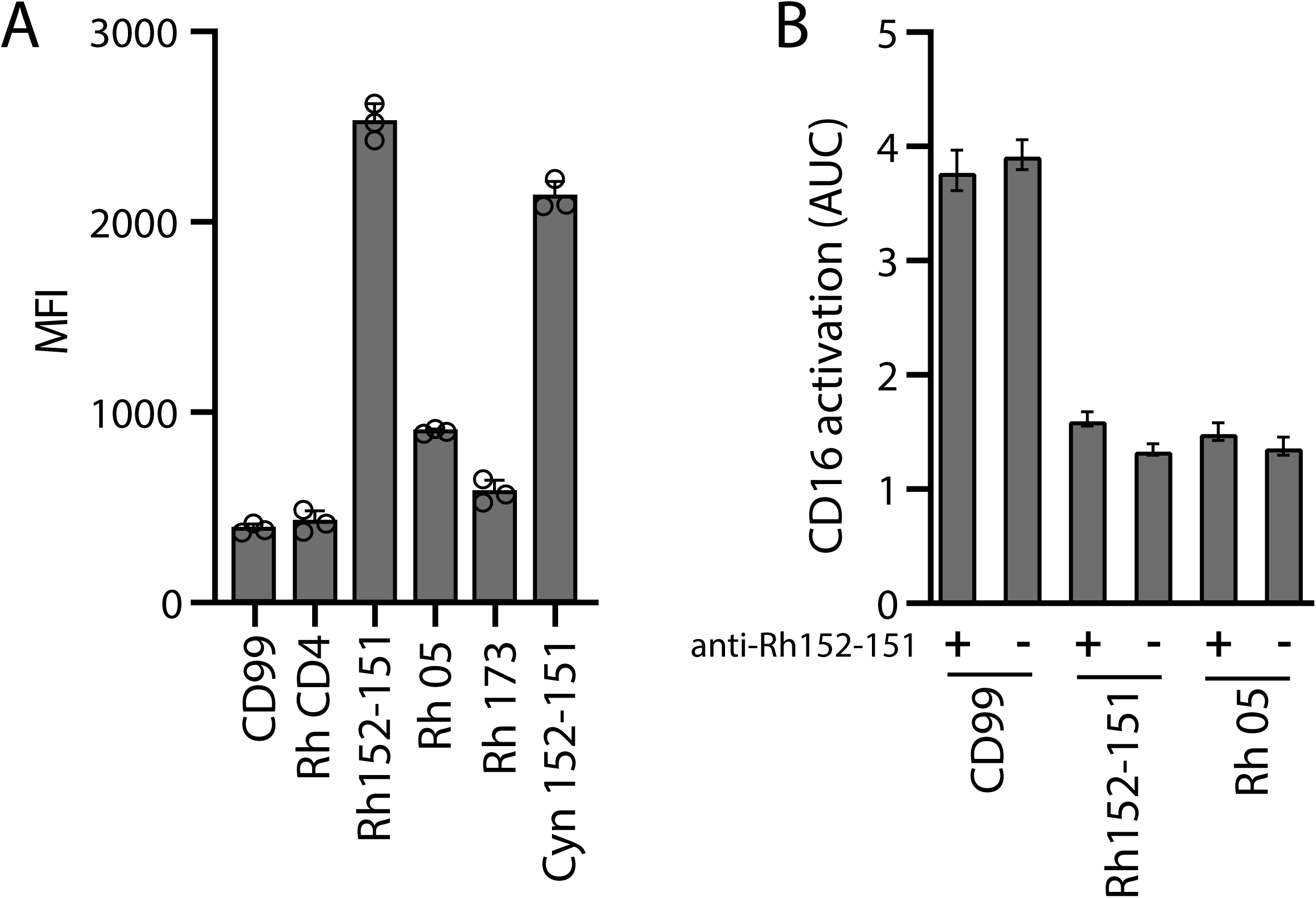
A Rh152/151 specific mAb from mouse immunization does not impact vFcγR function. A) Hela cells transfected with RhCMV vFcγRs or control proteins (rhesus-CD4, human CD99) were probed for recognition by the PE-labeld Rh152-151 specific mAb via flow cytometry. Cyn = Cynomolgus. Rh = Rhesus. Three independent experiments. Error bars = +SD. B) Human CD16 activation was tested on Hela cells transfected with rhesus-CD4 and the indicated vFcγRs or a human CD99 control from T2A-linked constructs to ensure equimolar expression. Cells were pre-incubated with the Rh152-151 specific mAb or not before addition of rhesus-CD4 specific mAb and co-culture with human CD16 reporter cells. Three independent experiments. Error bars = +SD.

**Fig S4.**
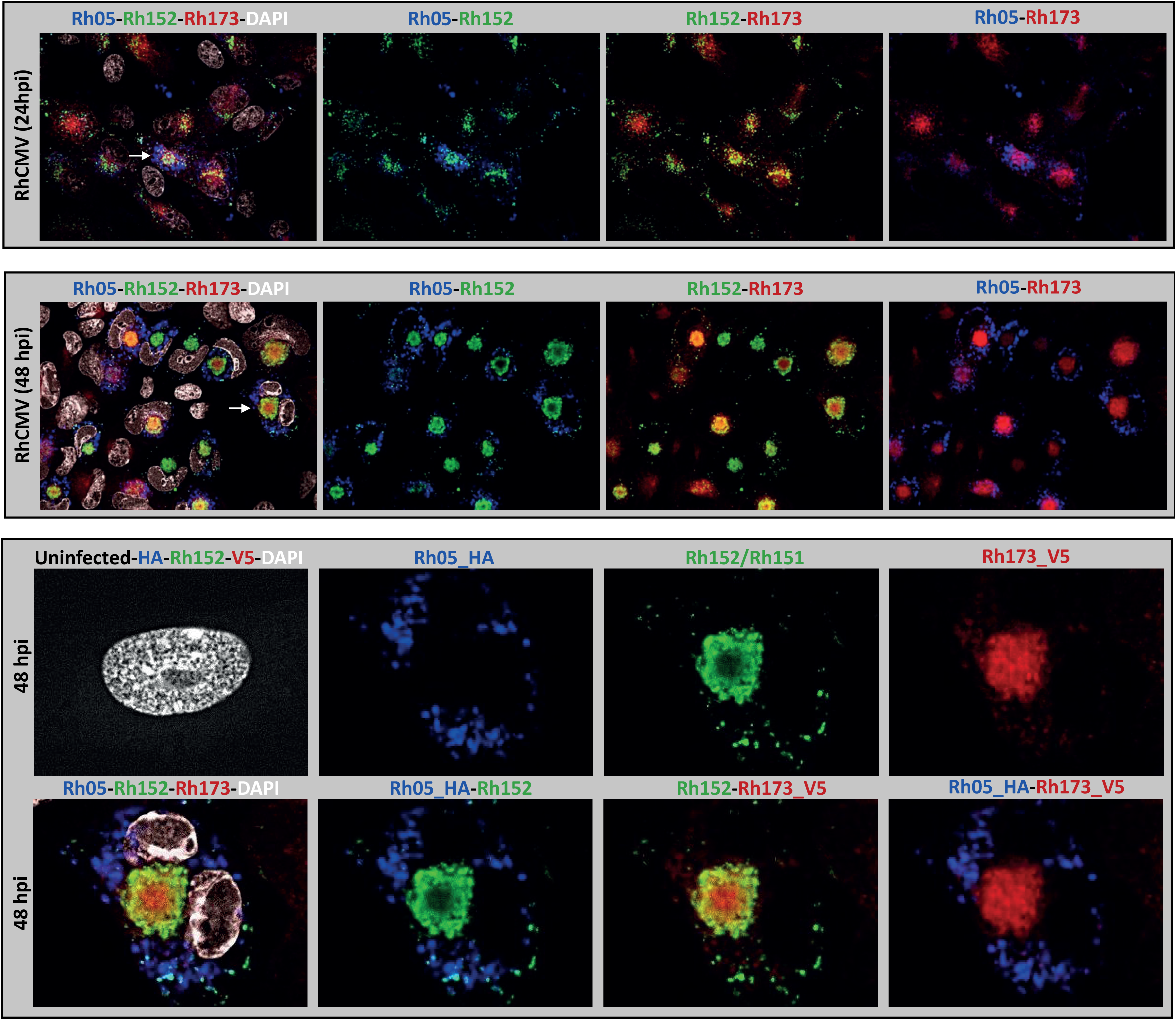
IFA for vFcγR co-localization at 24h post-infection. tRFs were infected with RhCMV FL+Rh05-HA+Rh173-V5 at MOI 1 and immunofluorescence assay was performed at 24 hpi and 48 hpi. Cells were probed using anti-HA, anti-Rh152/151 and anti-V5 antibodies for detecting vFcγRs. Data presented in the top row is a 100X microscopic view image at 24 hpi (cropped images in Fig 3F), while the center row shows the 100X microscopic view image at 48 hpi. The bottom row shows a cropped representative single cell at 48 hpi (marked with white arrow).

**Fig S5.**
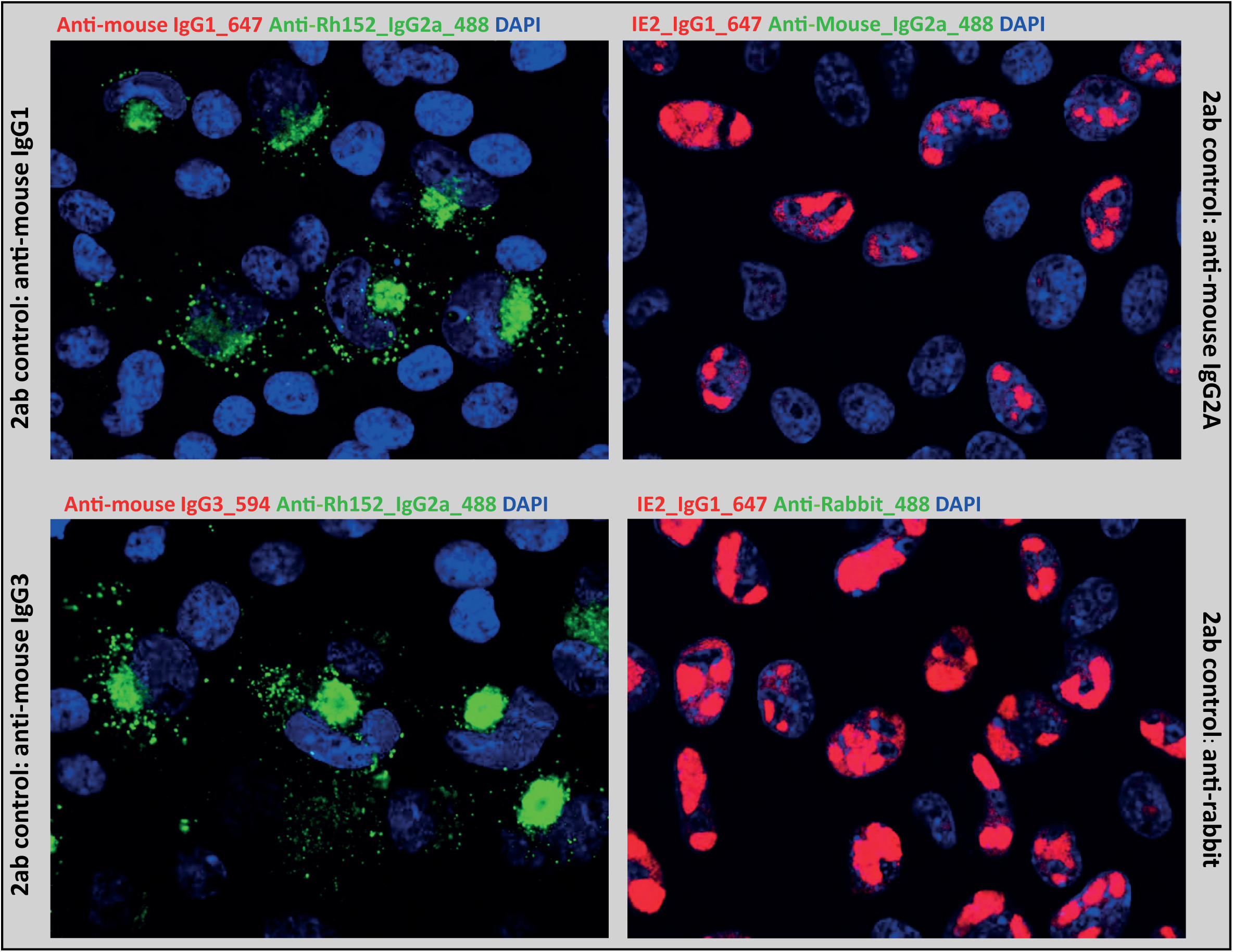
Control IFA for secondary antibody binding. To investigate whether secondary antibodies could bind to vFcγRs and result in non-specific signals, tRFs were infected with FL-RhCMV/Rh05-HA/Rh173-V5 at MOI 1 and IFA was performed at 48 hpi. Cells were probed using either anti-IE2 or anti-RhCMV antibodies for detecting infected cells, and additionally also probed with all florescent conjugated secondary antibodies. No non-specific vFcγR binding of secondary antibodies was observed in infected cells.

**Fig S6.**
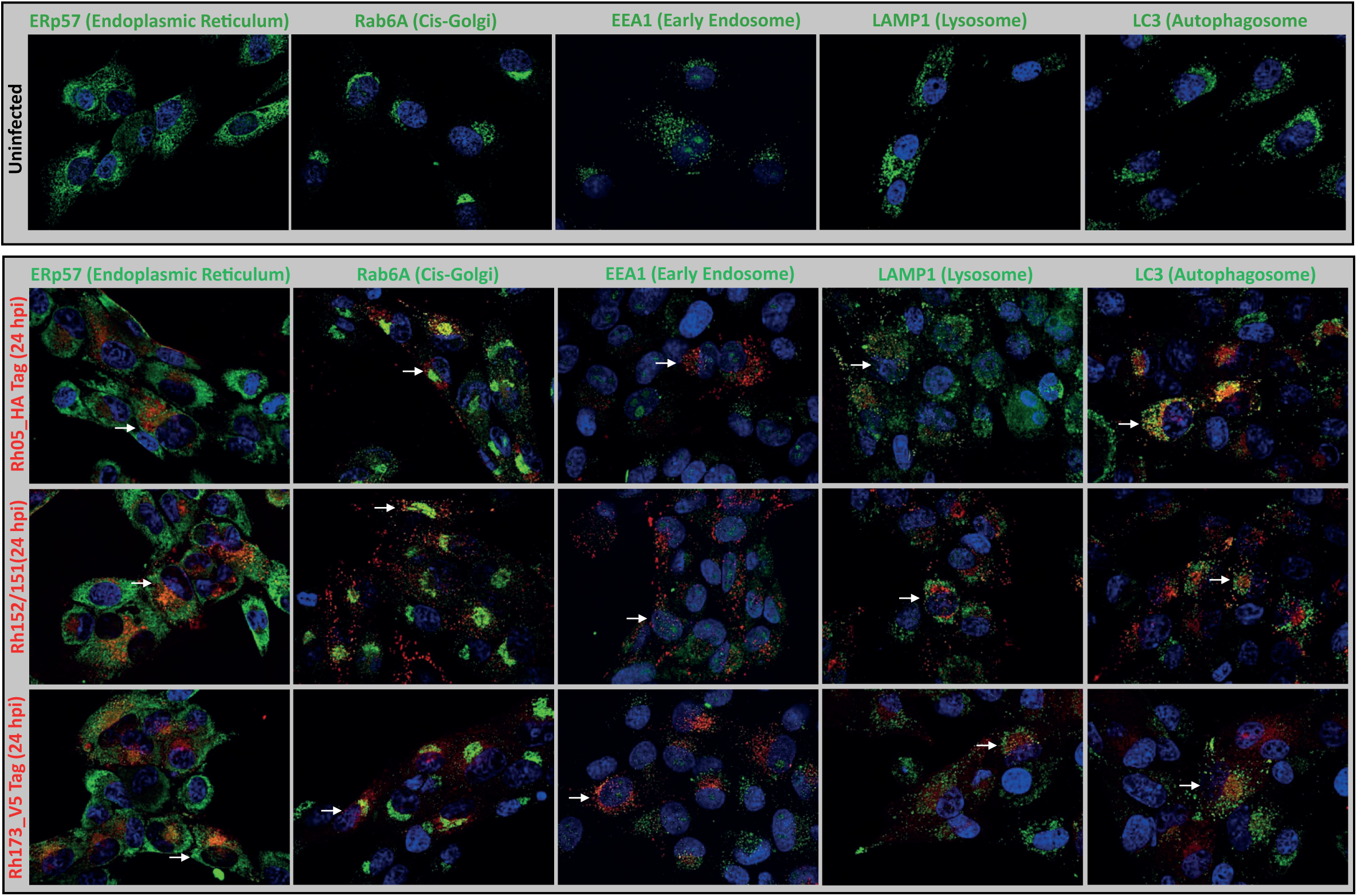
Co-localization of vFcγRs with markers of cellular organelles. Organelle staining pattern in uninfected tRF cells are shown in top row. In bottom row, tRFs were infected with FL-RhCMV/Rh05-HA/Rh173-V5 at MOI 1 and IFA was performed at 24 hpi. Cells were probed using anti-HA, anti-Rh152/151 and anti-V5 antibodies for detecting vFcRs. Additionally, cells were probed using anti-ERp57, Rab6A, LAMP1 and LC3 for detecting specific organelles. Data presented in bottom row is the complete 100X microscopic view image at 24 hpi (cropped images in Fig 3G).

**Fig S7.**
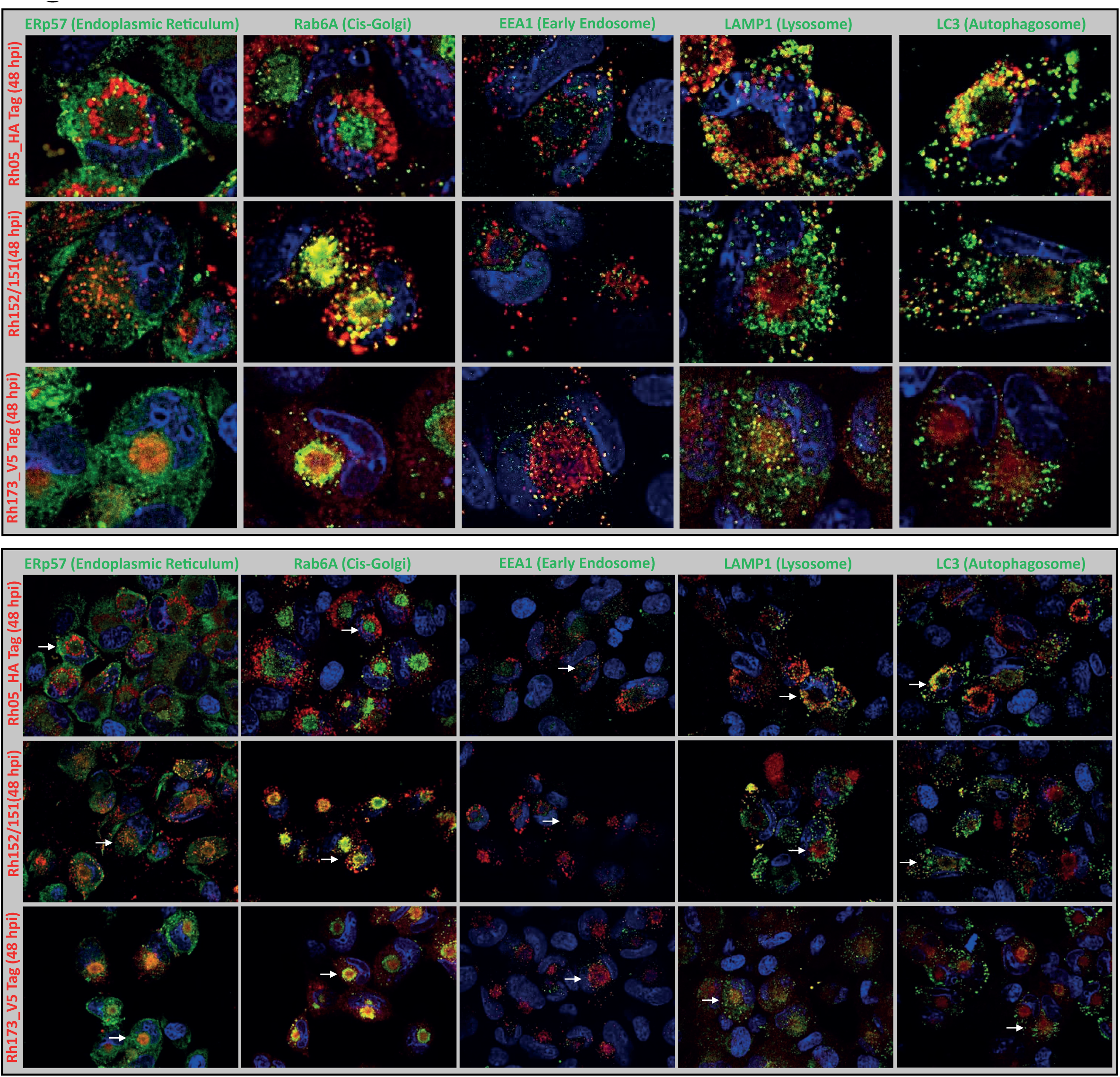
IFA for vFcγR co-localization at 24h post-infection. tRFs were infected with FL-RhCMV/Rh05-HA/Rh173-V5 at MOI 1 and immunofluorescence assay was performed at 48 hpi. Cells were probed using anti-HA, anti-Rh152/151 and anti-V5 antibodies for detecting vFcRs. Additionally, cells were probed using anti-ERp57, Rab6A, LAMP1 and LC3 for detecting specific organelles. The data presented in top panel is cropped representative single cell view from 100X microscopic view image. Bottom row shows cropped representative single cell at 48 hpi (marked with white arrow).

